# The role of LEDGF in transcription is exploited by HIV-1 to position integration

**DOI:** 10.1101/2024.06.29.601340

**Authors:** Rakesh Pathak, Caroline Esnault, Rajalingam Radhakrishnan, Parmit K Singh, Hongen Zhang, Ryan Dale, Abhishek Anand, Gregory J Bedwell, Alan N Engelman, Ali Rabi, Sahand Hormoz, Priyanka Singh, Henry L Levin

## Abstract

HIV-1 integration occurs across actively transcribed genes due to the interaction of integrase with host chromatin factor LEDGF. Although LEDGF was originally isolated as a co-activator that stimulates promoter activity in purified systems, this role is inconsistent with LEDGF-mediated integration across gene bodies and with data indicating LEDGF is a histone chaperone that promotes transcriptional elongation. We found LEDGF is enriched in pronounced peaks that match the enrichments of H3K4me3 and RNA Pol II at transcription start sites (TSSs) of active promoters. Our genome-wide chromatin mapping revealed that MLL1 had a dominant role in recruiting LEDGF to promoters and the presence of LEDGF recruits RNA Pol II. Enrichment of LEDGF at TSSs correlates strongly with levels of integration across the transcribed sequences, indicating that LEDGF at TSSs contributed to integration across gene bodies. Although the N-terminal Pro-Trp-Trp-Pro (PWWP) domain of LEDGF interacts with nucleosomes containing H3K36me3, a modification thought to recruit LEDGF to chromatin, we found H3K36me3 does not contribute to gene specificity of integration. These data support a dual role model of LEDGF where it is tethered to promoters by MLL1 and recruits RNA Pol II. Subsequently, LEDGF travels across genes to effect HIV-1 integration. Our data also provides a mechanistic context for the contribution made by LEDGF to MLL1-based infant acute leukemia and acute myeloid leukemia in adults.

## Introduction

The integration of viral DNA (vDNA) into chromosomes is an essential and irreversible feature of HIV-1 infection that eradicates the immune system and establishes a latent reservoir of T cells with intact proviruses ^1–3^. Much has been learned about the transport of the pre-integration complex (PIC) through nuclear pores and the subsequent role of CPSF6 in moving the PIC from the periphery into the nuclear lumen ^4–7^. The PIC associates with nuclear speckles and the chromatin associated transcription factor LEDGF/p75 interacts directly with integrase (IN), causing integration to occur across the bodies of actively transcribed genes ^8–14^. The integration positioned by LEDGF is in gene dense regions in proximity to nuclear speckles ^11,15^. LEDGF has modular domains that support a range of DNA and protein interactions (Fig. 1A). The N-terminal Pro-Trp-Trp-Pro (PWWP) domain that interacts with histone H3K36me3 modified nucleosomes and the DNA binding AT-hooks that promote chromatin binding are common to both the long p75 and short p52 isoforms of LEDGF. The C-terminal integrase binding domain (IBD) that associates tightly with lentivirus integrase (IN) proteins as well as a number of other proteins including transcription factors IWS1, MED1, and MLL1 ^16–22^ is unique to the p75 isoform. Here, we use LEDGF to refer to LEDGF/p75.

**Fig. 1.**
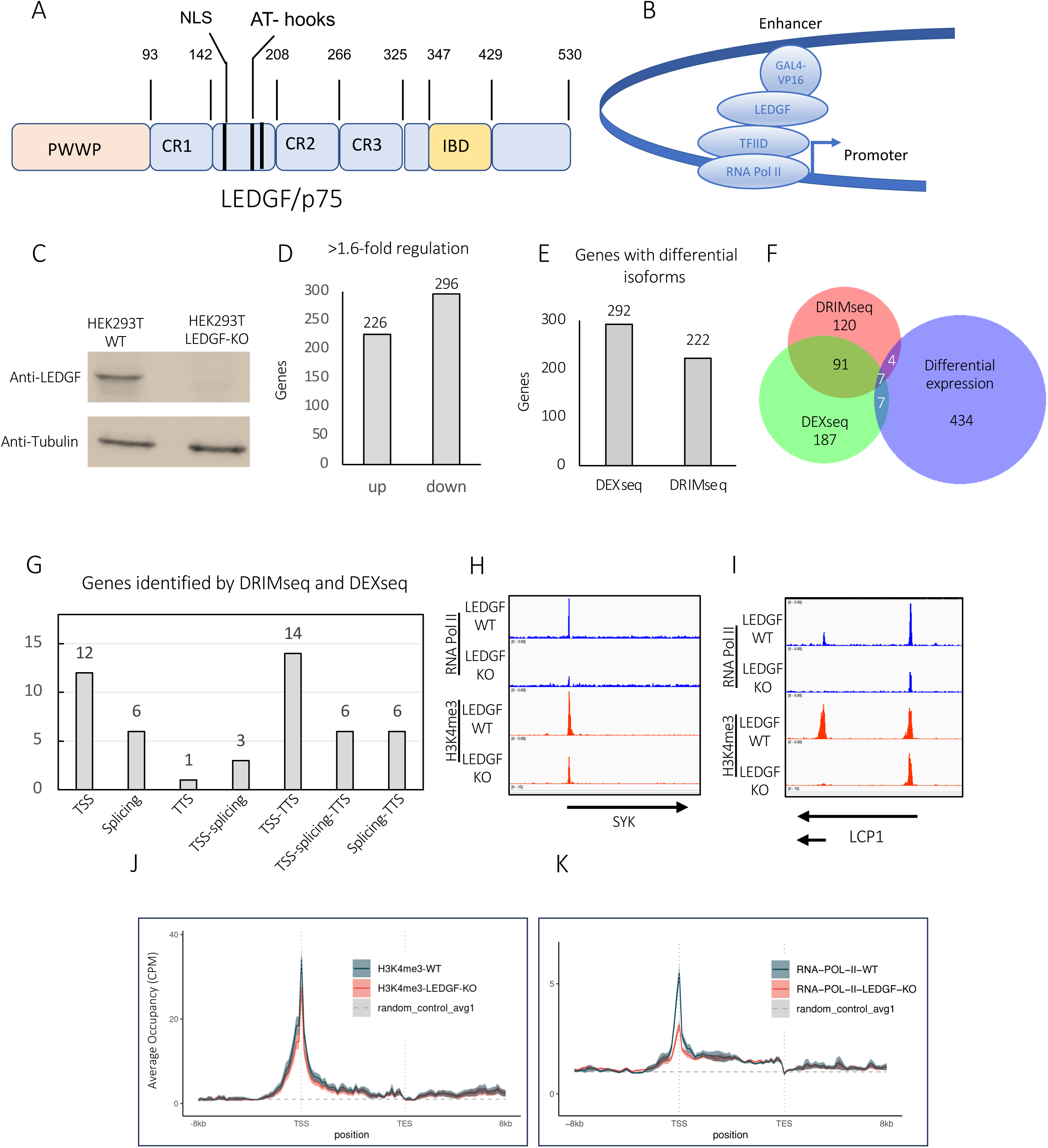
LEDGF regulates gene expression. A. The domains of LEDGF include the N-terminal Pro-Trp-Trp-Pro (PWWP), charge regions (CRs) 1-3, nuclear localization sequence (NLS), AT-hooks, and the integrase binding domain (IBD). B. Model depicting the co-activator activity of LEDGF in biochemical assays. C. Immunoblot of WT and LEDGF KO cells. D. RNA-seq results of LEDGF KO cells showing genes differentially expressed >1.6-fold. DEX-seq and DRIM-seq results of LEDGF KO cells showing genes with differential transcript usage. F. Overlap between genes with differential expression and genes with differential transcript usage. G. Number of genes with differential usage transcripts with different transcript start sites (TSS), altered splice choice (splicing), and different transcript termination sites (TTS). H and I. Chip-Seq enrichments of RNA Pol II and HeK4me3 in WT and LEDGF KO cells. J. and K. Metagene analyses of RNA Pol II and H3K4me3 in genes down regulated >1.6-fold in LEDGF KO cells. The random control shows the enrichment expected for random association across the genome (dashed line).

The chromatin at sites of HIV-1 integration is enriched with various histone modifications present during active transcription including H4K16ac, H3K36me3, and H3K4me1 ^11^. H3K36me3 has the potential to play a direct role in integration as the PWWP domain of LEDGF is shown in biochemical and structural studies to specifically associate with H3K36me3 modified nucleosomes ^20,23–25^. Cells lacking LEDGF do not exhibit integration across gene bodies but instead show a marked preference for sites near the start of transcription ^12,13,26,27^.

In addition to distributing integration across gene bodies, LEDGF interacts with splicing machinery and has histone chaperone activity that can promote RNA Pol II elongation ^19,27,28^. However, these activities of LEDGF across gene sequences are at odds with the original discovery of LEDGF that showed in purified systems the factor binds promoters and functions as a transcriptional coactivator ^29^. While LEDGF may possess all these activities, direct evidence of each requires high specificity mapping of positions where LEDGF binds chromatin. Current information about LEDGF association with chromatin is limited. The lack of information about the gene specificity of its chromatin association also limits mechanistic understanding about why LEDGF is central to the cancer-causing effects of MLL1 rearrangements resulting in fusion proteins that occur in greater than 70% of infant acute leukemia and 10% of acute myeloid leukemia (AML) cases in adults ^30–32^.

Here, we have developed reagents that allow for the highly specific positioning of LEDGF that reveal an association with RNA Pol II at active promoters. With these tools we mapped promoter binding activity of LEDGF to an association of the IBD with MLL1. Integration levels across genes correlated with peaks of LEDGF at transcription start sites (TSSs). In addition, we found that the PWWP domain does not contribute significantly to the binding of LEDGF to RNA Pol II promoters and that H3K36me3 contributes modestly to the distribution of HIV-1 integration across transcribed sequences but not to the gene specific frequencies of integration.

## Results

LEDGF was originally identified as a transcription co-activator, bridging interactions between activators such as VP16 and general transcription machinery (Fig. 1B) ^29^. Its discovery as a cofactor in HIV-1 integration resulted in an extensive period of research that has all but eclipsed further studies of the role LEDGF plays in transcription. We sought to further characterize the cellular contributions of LEDGF to transcription because this knowledge has the potential to identify mechanistic features of HIV-1 integration not previously appreciated.

We conducted RNA-seq studies of HEK293T cells edited with CRISPR to lack LEDGF expression beyond amino acid 26 (Fig. 1C). Differential expression analysis showed that the absence of LEDGF in the knockout (KO) cells resulted in increased expression of 226 genes and reduced expression of 296 genes when applying a threshold of greater than 1.6-fold change (Fig. 1D). Most of the genes that are differentially expressed do not change more than 4-fold (Suppl. Figs. S1 and S2). The interaction of LEDGF with splicing factors ^19,27^ led us to ask whether the absence of LEDGF results in changes to alternative splicing. We employed two independent programs that compare RNA-Seq datasets to identify genes with differentially expressed spliced isoforms. Changes in isoform usage of an individual gene upon LEDGF depletion can be the result of altered splicing. The DEX-seq and DRIM-seq pipelines identified 292 and 222 genes that exhibited differentially expressed mRNA isoforms, respectively (Fig. 1E). Among these two sets, 98 genes were shown by both pipelines to have differentially expressed isoforms (Fig. 1F). Of these 98 examples we evaluated the interpretable transcript models of 48 genes with discordant changes in isoforms. We examined the isoform models for genes with changes in splice choices, TSSs, and transcription termination sites (TTSs) (Suppl. Fig. S3). While we found 21 of 48 genes had differential levels of mRNA isoforms with altered splice patterns, 9 of these had different TSSs and 12 had altered TTSs (Fig. 1G). Just 6 of 48 genes had only changes in splice sites, leaving open the possibility that LEDGF could be causing differential expression of most isoforms by changing levels of transcription initiation (TSS) or termination (TTS). A model that LEDGF contributes primarily to promoter activity is consistent with the result that 35 of the 48 genes identified by DRIM-seq and DEX-seq had pairs of differentially expressed isoforms with different TSSs. Importantly, just 18 of the 434 genes differentially regulated in the absence of LEDGF were identified by either DRIM-seq or DEX-seq, indicating that in terms of the number of genes effected, any role LEDGF plays in alternative splicing is minor relative to its function in promoter activity (Fig. 1F).

We evaluated the role of LEDGF at promoters that are down regulated in LEDGF KO cells by measuring their levels of RNA Pol II and H3K4me3, a histone modification closely associated with active promoters ^33–36^. ChIP-seq data at some down regulated genes showed reduced RNA Pol II and H3K4me3 in the LEDGF KO cells (Figs 1H and 1I). Metagene profiles of all 296 down regulated genes shows modest reduction in H3K4me3 and a substantial decrease in RNA Pol II at the TSSs (Figs. 1J and 1K).

### LEDGF is enriched at TSSs and recruits RNA Pol II

To test whether LEDGF directly contributes to promoter activity and as a consequence steady state levels of mRNA, we would need to compare amounts of LEDGF to levels of H3K4me3 and RNA Pol II at individual promoters. However, the availability of highly specific and dense chromatin-immunoprecipitation-DNA sequencing (ChIP-seq) data for LEDGF is limited due to the lack of antibodies that are effective in chromatin immunoprecipitation. We overcame this challenge with CRIPSR by inserting a biallelic 3XFLAG tag at the C-terminus of LEDGF in its native gene PSIP1 in HEK293T cells. With anti-FLAG antibodies we obtained ChIP-Seq data for LEDGF and compared it to our ChIP-seq of RNA Pol II and H3K4me3. The IP results of the cells expressing LEDGF-3XFLAG relative to cells lacking the 3XFLAG tag revealed that many genes had narrow peaks of LEDGF enrichment that corresponded closely with the promoter bound RNA Pol II and H3K4me3 (Figs. 2A and 2B). A full 95% and 54% of LEDGF-3XFLAG peaks (>2.5-fold enrichment) overlapped with H3K4me3 and RNA Pol II enrichment, respectively (MACS2, FDR<0.05). Thirteen percent of RNA Pol II and 11% of H3K4me3 peaks overlap with LEDGF-3XFLAG, suggesting that LEDGF occupies a subset of promoters. LCP1 is an example of a down regulated gene in the absence of LEDGF. It possesses promoter peaks of LEDGF that overlapped RNA Pol II and H3K4me3 enrichment which was reduced in the cells lacking LEDGF (Fig. 2A). SYK was also down regulated in cells lacking LEDGF and had promoter bound LEDGF with enrichment of RNA Pol II that was significantly reduced in the LEDGF KO cells. Genome-wide metagene profiles of LEDGF-3XFLAG had maximum enrichment at TSSs that mimic profiles of H3K4me3 and RNA Pol II (Figs 2C, 2D, and 2E). Profiles of genes with >2.5-fold enrichment of LEDGF-3XFLAG had much higher levels of H3K4me3 and RNA Pol II at TSSs indicating that LEDGF has a strong association with transcriptionally active promoters (Figs. 2F and 2G). We evaluated the genome-wide contribution of LEDGF at promoters with metagene plots of H3K4me3 and RNA Pol II. While the enrichment of H3K4me3 at TSSs was similar in cells lacking LEDGF to wild type cells, amounts of RNA Pol II were reduced in LEDGF KO cells (Figs. 2D and 2E). The contribution of LEDGF to RNA Pol II recruitment was most pronounced at genes with the highest enrichment of LEDGF as seen in metagene profiles of genes separated into five quantiles sorted by levels of LEDGF-3XFLAG (Suppl. Fig. S4). Of all RNA Pol II peaks that overlapped with LEDGF-3XFLAG, 93% were significantly reduced in absence of LEDGF (Diffbind, FDR<0.05). These data indicate that LEDGF is enriched at the TSSs of genes where it recruits RNA Pol II.

**Fig. 2.**
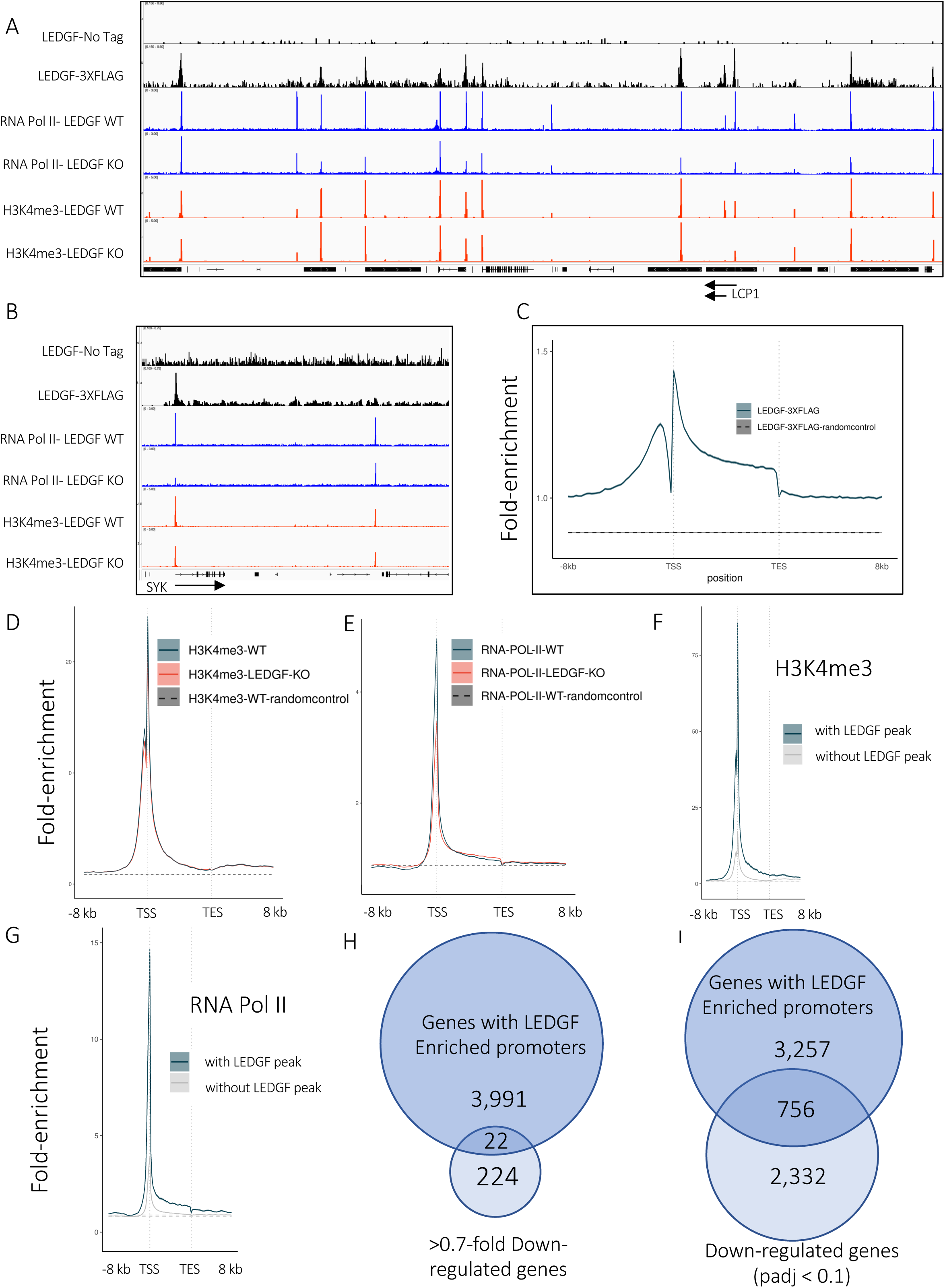
LEDGF is enriched at TSSs and recruits RNA Pol II. A. and B. ChIP-seq enrichment data of LEDGF-3XFLAG, RNA Pol II, and HeK4me3 from WT and LEDGF KO cells. C. D. and E. Metagene plots of fold-enrichment of LEDGF-3XFLAG (C) H3K4me3 (D) and RNA Pol II (E) in WT and LEDGF KO cells. The dashed line is a control indicating random enrichment across the genome. F. and G. Metagene plots of fold-enrichment H3K4me3 (F) and RNA Pol II (G) of genes with and without peaks of LEDGF-3XFLAG. H. and I. Overlap of genes with LEDGF at promoters and genes downregulated in LEDGF KO cells by >0.7-fold (H) and with p adjusted <0.1 (I).

Although 4,013 genes have promoters bound by LEDGF-3XFLAG (>2.5X enrichment), just 22 of these were down regulated (>01.6-fold) in the LEDGF KO cells (Fig. 2H). While this was not a significant overlap, if considering the broader group of all genes downregulated (padj < 0.1, odds ratio 1.08, Fisher’s exact test) in LEDGF KO cells a larger fraction had LEDGF bound at the TSS (756/2,332) which does represent a significant overlap (p < 2.2e-16, odds ratio 4.16, Fisher’s exact test, Fig. 2I). The relatively small overlap of genes substantially dependent on LEDGF for expression and genes with LEDGF enriched at TSSs suggests that the function LEDGF has at most promoters is redundant with other transcription factors, insufficiently active to increase transcription despite changing the amount of Pol II, or the steady-state RNA levels we measure are indirectly modulated post-transcriptionally.

### LEDGF is recruited to TSSs of genes by MLL1

LEDGF associates with nucleosomes containing H3K36me3 due to a hydrophobic cavity in the PWWP domain that interacts with the trimethylated lys36 ^20,23,24^. Because the evidence for this interaction is largely from biochemically purified systems, much less is known about the importance of H3K36me3 in the *in vivo* specificity of LEDGF binding to chromatin. For example, H3K36me3 is typically enriched towards the 3’ regions of genes (Fig. 3A, left panel) ^37^. This distribution is at odds with the profile of LEDGF dependent integration which is higher near the 5’ end of gene bodies as observed in cultured cells (Fig. 3A, center and right panels) ^13,27^. In particular, the positions of LEDGF-3XFLAG at TSSs (Figs. 2A, 2B, and 2C) varies greatly from the patterns of H3K36me3 (Figs. 3A and 3B). While the profile of 1 million insertions described by Singh et al does show integration across transcribed sequences of individual genes, the regions with highest integration do not correspond with the highest levels of H3K36me3 (Fig. 3B) ^27^.

**Fig. 3.**
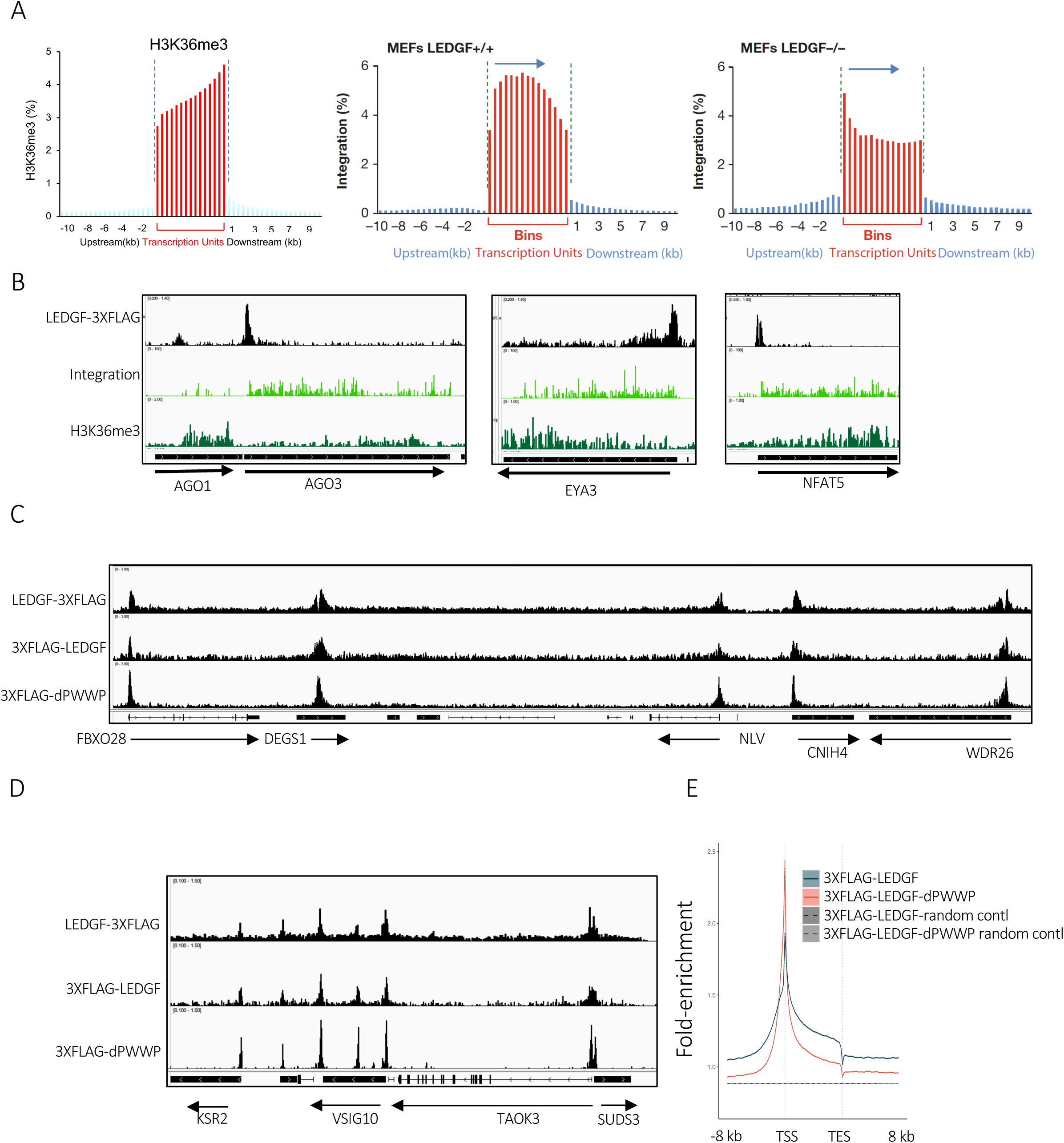
The recognition of H3K36me3 by the PWWP domain does not position LEDGF at TSSs. A. Metagene plots of H3K36me3 and integration in HEK293T cells and MEFs, respectively. Integration in MEFs was reproduced from Singh et al ^27^. B. Examples of genes with LEDGF-3XFLAG at TSSs showing integration and H3K36me3. C. and D. Examples of genes with ChIP-seq enrichment of LEDGF-3XFLAG, 3XFLAG-LEDGF, and 3XFLAG-LEDGF-dPWWP. E. Metagene plot of genes with LEDGF enrichment showing 3XFLAG-LEDGF and 3XFLAG-LEDGF-dPWWP.

We tested the role of the PWWP domain in the chromatin association of LEDGF by ectopically expressing an N-terminally tagged 3XFLAG-LEDGF in LEDGF KO HEK293T cells. The 3XFLAG-LEDGF expressed from an integrated Sleeping Beauty vector was enriched at TSSs with a profile similar to the endogenously expressed LEDGF-3XFLAG (Figs. 3C and 3D) with 98% of the 7,161 LEDGF-3XFLAG peaks overlapping peaks of 3XFLAG-LEDGF (MACS2, FDR<0.05, run 11). Due to significantly higher number of called peaks in the 3XFLAG-LEDGF data (260,991), 1.7% overlapped with LEDGF-3XFLAG. Despite the visual similarity between these two datasets, a higher level of noise in this 3XFLAG-LEDGF dataset likely resulted in the higher number of called peaks.

The visual similarity in the enrichment profiles suggests that the position of the 3XFLAG tag does not alter the specificity of the LEDGF chromatin profiles. Importantly, the chromatin association of 3XFLAG-LEDGF lacking the PWWP domain (3XFLAG-dPWWP) and its expression was not significantly different from intact 3XFLAG-LEDGF (Figs. 3C, 3D and Suppl Fig. S5A). The genome-wide metagene profile of 3XFLAG-dPWWP retained the high enrichment at TSSs comparable to profile of wild type 3XFLAG-LEDGF (Fig. 3E). Interestingly, the removal of the PWWP domain resulted in higher enrichment at TSSs and lower levels in the body of genes. This redistribution of enrichment towards the TSSs indicates there is competition between LEDGF bound in transcribed sequences verses LEDGF at promoters and the PWWP domain contributes to the downstream pool, consistent with PWWP being required for interacting with H3K36me3 which is found more downstream.

An alternative domain of LEDGF that could be involved with its chromatin association at promoters is the IBD. A Sleeping Beauty vector encoding LEDGF with a C-terminal truncation (3XFLAG-LEDGF-1-325) lacked the IBD and expressed protein at levels similar to the full-length LEDGF (3XFLAG-LEDGF) (Suppl. Fig. S5B). Similar to 3XFLAG-LEDGF with IBD intact, enrichment of 3XFLAG-LEDGF-1-325 relative to input matched the peaks of RNA Pol II, and H3K4me3 at TSSs (Figs 4A, 4B, and 4C). Notably, LEDGF lacking the IBD (3XFLAG-LEDGF-1-325) had much less enrichment at the positions bound by the full-length LEDGF. Differential binding analysis showed that 81% of 3XFLAG-LEDGF enriched peaks were significantly reduced in the profile of 3XFLAG-LEDGF-1-325 (FDR<0.05). Genome-wide metagene profiles confirmed that 3XFLAG-LEDGF-1-325 associated substantially less with TSSs compared to the full-length 3XFLAG-LEDGF indicating that the IBD region of LEDGF plays a significant role in the association of LEDGF with promoters (Fig. 4D).

**Fig. 4.**
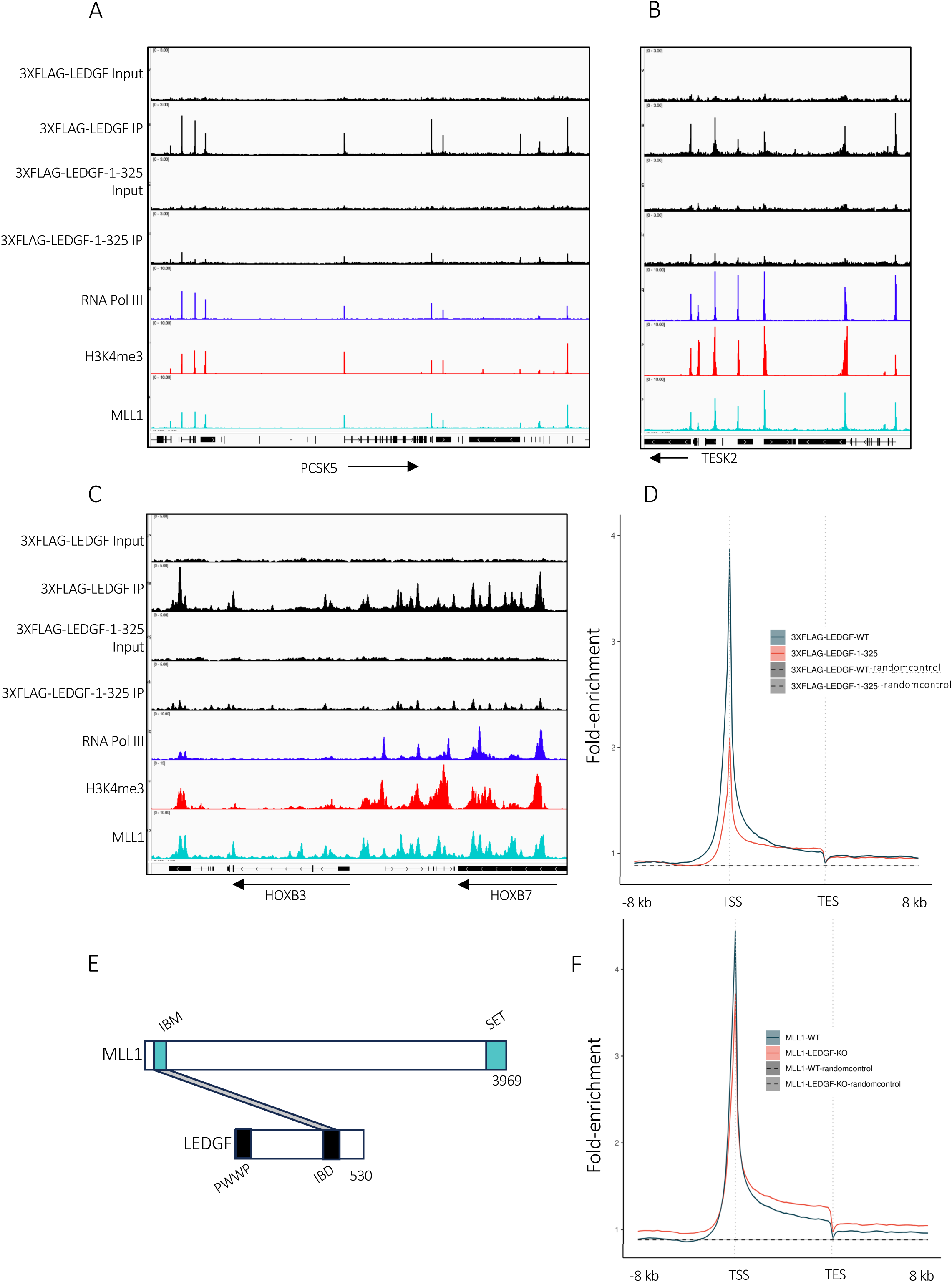
The IBD is required to recruit LEDGF to TSSs. A. B. and C. Examples of genes with enrichment of 3XFLAG-LEDGF, 3XFLAG-LEDGF-1-325, RNA Pol II, H3K4me3, and MLL1. D. Metagene plot showing fold enrichment of 3XFLAG-LEDGF and 3XFLAG-LEDGF-1-325. The dashed line is a control indicating random enrichment across the genome. E. Diagram of MLL1 and LEDGF indicating interaction between IBM and IBD. F. Metagene plot showing fold enrichment of MLL1 in WT and LEDGF KO cells.

The LEDGF IBD has been shown to form a structured complex with an intrinsically disordered module of MLL1, the IBD-binding motif (IBM) and menin (Fig. 4E) ^21,38^. By mapping chromatin association with ChIP-Seq we found MLL1 was enriched in narrow peaks that closely matched the profiles of RNA Pol II, H3K4me3, and 3XFLAG-LEDGF at TSSs (Figs. 4A and 4B). The close association of MLL1 with RNA Pol II and H3K4me3 extended across the HOX loci, a region strongly regulated by LEDGF and MLL1 (Fig. 4C) ^39–42^. Interestingly, the genome-wide profile of MLL1 across gene bodies was specifically enriched at TSSs and showed strong similarity to the enrichment of 3XFLAG-LEDGF (compare black lines in Figs 4D and 4F). Genome-wide, 47% of 3XFLAG-LEDGF enriched peaks >2.5-fold overlapped with sites of MLL1 with >2.5-fold enrichment (MACS2, FDR<0.05, qval1^-10^, run 9 data) and 85% of MLL1 peaks overlapped with 3XFLAG-LEDGF (MACS2, FDR<0.05, qval1^-10^, run 9). The overlap of LEDGF and MLL1 was also readily observed with ChIP-Seq data of LEDGF-3XFLAG with 54% of peaks overlapping with MLL1 and 12 % of the MLL1 sites overlapped with LEDGF-3XFLAG (>2.5-fold enrichment (MACS2, FDR<0.05, qval1^-10^) (Figs. 5A and 5B). The high level of overlap of MLL1 and LEDGF positions on chromatin raised the possibility that the IBD interaction with MLL1 tethers LEDGF to chromatin at the TSSs. We tested this using shRNA which greatly reduced levels of MLL1 and found this consequently diminished the chromatin association of LEDGF-3XFLAG (Figs. 5C and 5D). Whole cell levels of LEDGF-3XFLAG were not reduced (Fig. 5C). Genome-wide analysis of cells with reduced MLL1 showed 96% of LEDGF-3XFLAG peaks were significantly reduced (Diffbind, FDR<0.05 of enrichment q<1^-10^) particularly at TSSs bound by LEDGF (Figs. 5E and 5F). MLL1 is one of six mixed lineage leukemia (MLL) family methyltransferases responsible for methylation of histone H3K4 ^43,44^. However, the shRNA knockdown of MLL1 indicates that at least at TSSs bound by LEDGF there was little change in levels of H3k4me3 (Fig. 5G). In addition to the role MLL1 plays in the association of LEDGF at TSSs, there was a reciprocal relationship where MLL1 enrichment at TSSs was enhanced by LEDGF (Fig. 4F). Sixty-nine percent of MLL1 peaks that overlapped LEDGF were significantly reduced in LEDGF KO cells (Diffbind, FDR<0.05).

**Fig. 5.**
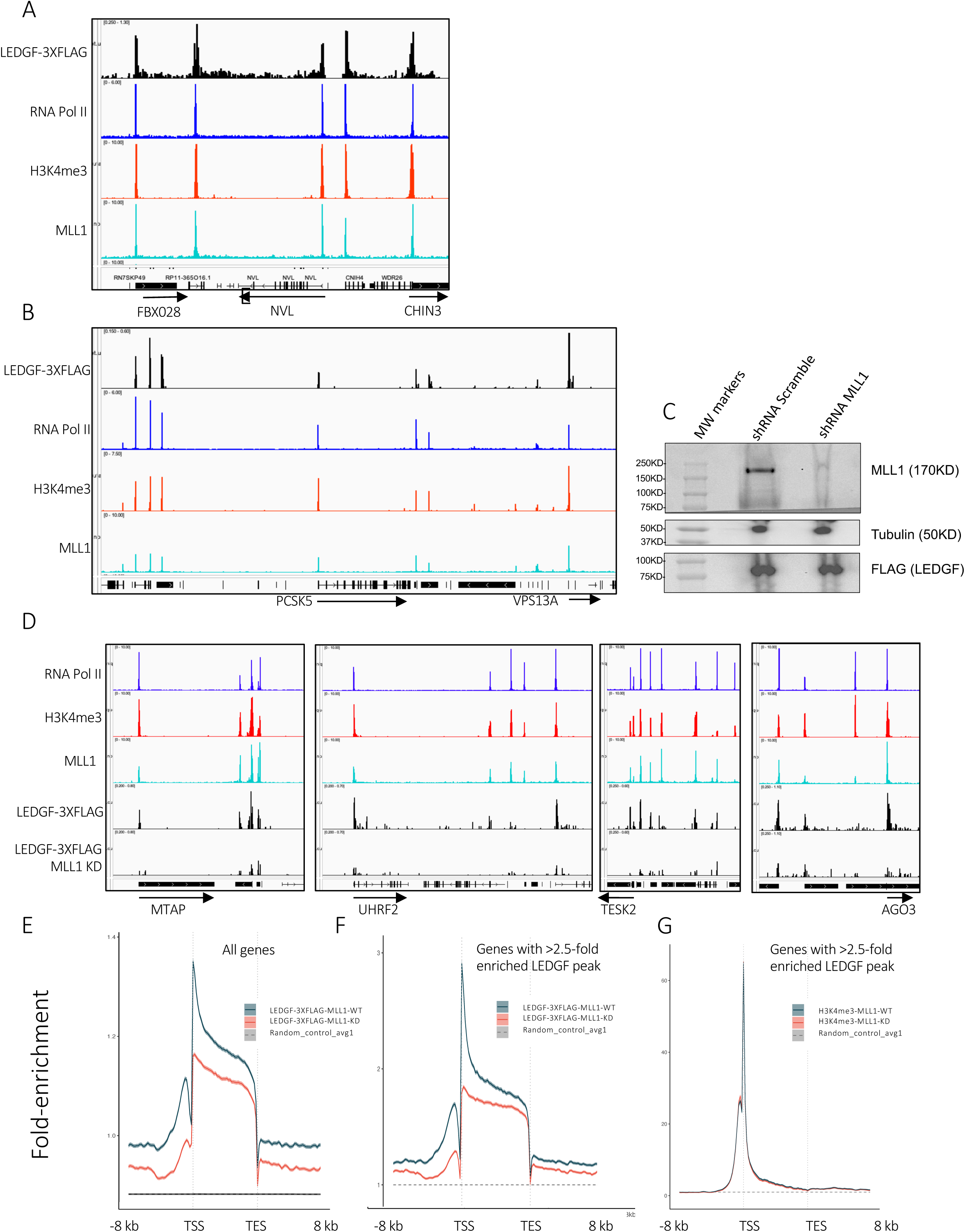
MLL1 recruits LEDGF to TSSs. A. and B. Examples of genes with enrichment of LEDGF-3XFLAG, RNA Pol II, H3K4me3, and MLL1. C. Immunoplot of cells expressing shRNA scramble or shRNA MLL1. D. Examples of genes with enrichment of LEDGF-3XFLAG in cells expressing shRNA scramble or shRNA MLL1. E. Metagene plots of fold-enrichment showing LEDGF-3XFLAG in cells expressing shRNA scramble or shRNA MLL1. F. and G. Metagene plots showing fold-enrichment of LEDGF-3XFLAG and H3K4me3 at genes with >2.5-fold enrichment of LEDGF from cells expressing shRNA scramble or shRNA MLL1. The dashed line is a control indicating random enrichment across the genome.

LEDGF-3XFLAG enriched at TSSs participates in integration of HIV-1 across gene bodies LEDGF plays a key role in HIV-1 infection by interacting with IN and positioning the vast majority of integration events throughout the sequences of active genes ^10,12,27,45^. The enrichment of LEDGF at TSSs is unexpected because this promoter region is upstream of where the majority of insertions occur. This discordance suggests the LEDGF bound at TSSs is independent of the pool of LEDGF responsible for integration or that this pool at the TSSs is due to the 3XFLAG tag. A contribution of the tag to TSS association seems unlikely because the N-terminal tagged LEDGF had an equivalent recognition of promoters as did the C-terminal tagged protein (Figs. 3C and 3D). In addition, we tested whether the 3XFLAG tag might have altered the function of LEDGF in integration. The integration mediated by LEDGF-3XFLAG had specificities that matched native LEDGF including the frequencies of insertions in genes, speckle associated domains (SPADs), lamin associated domains (LADs), TSSs, and gene dense regions (GDs) (Suppl. Fig. S6, compare KMT2A control to WT HEK297T or SETD2 control) ^1,5,15^. From this analysis we concluded that the C-terminal tag does not significantly alter integration specificity.

To evaluate whether the LEDGF at TSSs is involved in integration we tested the relationship between TSS bound LEDGF and integration by comparing the average integration density/kb in the genes with LEDGF peaks to those lacking LEDGF. Using HEK293T cell insertion data from Singh et al there was clearly higher integration densities in the genes with LEDGF-3XFLAG bound at TSSs (Suppl. Fig S7) ^27^. In an analysis of genes ranked by integration frequency and placed into bins of 100, there was a strong positive correlation with integration frequency and the number of LEDGF-3XFLAG peaks per bin (Fig. 6A). Interestingly, when numbers of integration sites in individual genes was compared to fold-enrichment of LEDGF-3XFLAG, we observed a threshold level of LEDGF binding that was required for efficient integration (Fig. 6B). No relationship was observed when LEDGF binding was compared to genes with randomly positioned integration sites. The correlation of LEDGF enrichment with integration levels indicates the LEDGF at TSSs participates in integration. One exception to the link between LEDGF association and integration are the HOX genes, where there is strong enrichment of LEDGF, MLL1, RNA Pol II, and H3K4me3 across the genes, but where there is little integration (Suppl. Fig. S8).

**Fig. 6.**
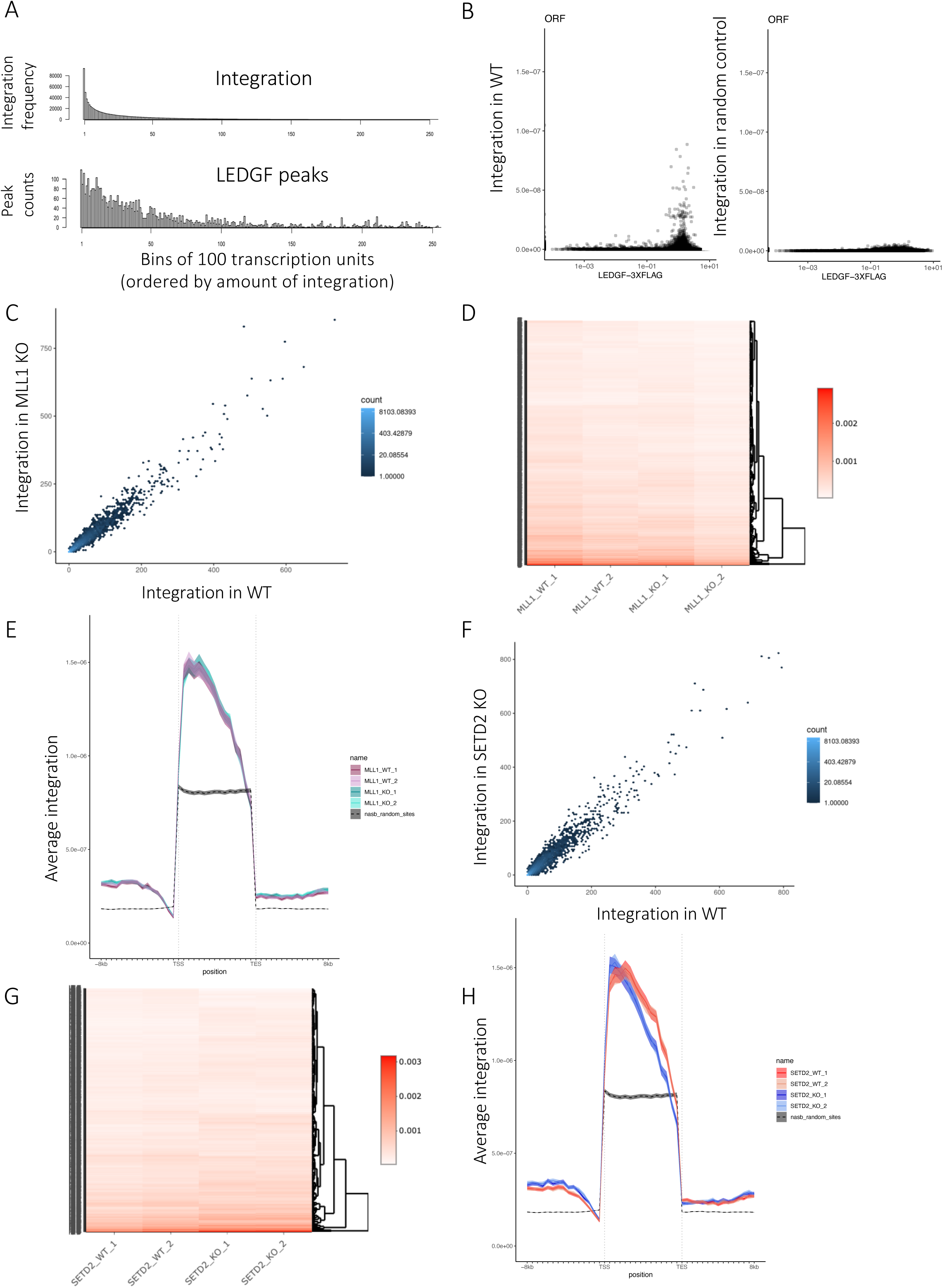
Integration profiles are largely unchanged in cells with MLL1 KD and cells lacking H3K36me3. A. Numbers of LEDGF peaks in genes (bottom panel) sorted by integration frequency (top panel) and binned into groups of 100. B. Left panel; Individual genes in scatter plot of fold-enrichment of LEDGF-3XFLAG vs integration frequency. Right panel; Individual genes in scatter plot of fold-enrichment of LEDGF-3XFLAG vs frequency of random integration sites. C. Integration frequency in individual genes mapped in WT and MLL1 KD cells. D. Heatmap of the 1000 genes with the highest integration frequency in cells expressing the shRNA scramble or shRNA MLL1 KD. E. Metagene plots of average integration in cells expressing the shRNA scramble or shRNA MLL1 KD. The dashed line is a control indicating random enrichment across the genome. F. Integration frequency in individual genes mapped in WT or SETD2 KO cells. G. Heatmap of the 1000 genes with the highest integration frequency in WT or SETD2 KO cells. H. Metagene plots of average integration in WT or SETD2 KO cells.

If the LEDGF enriched at TSSs is involved in integration, we might expect the cells with reduced MLL1 and lower LEDGF at TSSs to exhibit integration patterns similar to LEDGF KO cells. These behaviors are observed when LEDGF expression is entirely eliminated ^10,12,45^. Integration in cells lacking LEDGF have increased positions near TSSs, reduced recognition of genes, increased insertions in LADs, lower integration in SPADs and reduced insertions in GDs. However, these LEDGF independent behaviors did not occur in the MLL1 (KMT2A) KD cells (Suppl. Fig S6, compare KMT2A KD, WT, SETD2 control to LEDGF KO). And a comparison of the normalized amount of integration per gene showed the MLL1 (KMT2A) KD did not significantly alter the integration preferences of genes (Fig. 6C). A heatmap of integration density of the top 1,000 gene targets showed the MLL1 KD caused little change in selection of these genes (Fig. 6D). In a parallel test of altered integration specificity, we evaluated the recurrent integration gene (RIGs) targets and found the integration in MLL1 KD cells compared to the shRNA control had a Szymkiewicz–Simpson overlap coefficient of 0.95 where 1.0 is full overlap (Suppl. Table S1)^15^. When we evaluated integration in recurrent avoided genes (RAGs), the overlap coefficient shRNA control to MLL1 KD was 0.93, indicating the KD of MLL1 did not alter levels of integration in genes regularly avoided by HIV-1 (Suppl. Table S1). Integration specificity was also evaluated regarding genome annotations defined by 3D locations in the nucleus termed SPIN (Spatial Positioning Inference of the Nuclear genome) ^15,46^. These regions include areas associated with speckles, lamina, and varying radial depths of the nucleus. SPIN analysis indicated that reduced expression of MLL1 did not significantly alter integration preferences for SPIN states (Suppl. Table S2). The metagene profile of integration was unchanged in cells with reduced MLL1 (Fig. 6E). These data taken together indicate that the KD of MLL1 did not significantly alter integration specificity or distribution across gene bodies. However, given that LEDGF expression must be entirely eliminated in order to observe changes in integration behavior, it is possible that the residual amounts of LEDGF at TSSs of the MLL1 KD cells was sufficient to position integration ^10,12,45^.

The cells expressing 3XFLAG-LEDGF-dPWWP showed that the PWWP domain did not significantly contribute to LEDGF enrichment at TSSs (Figs. 3C, 3D, and 3E). This result is unexpected because of reports that the PWWP domain is important for the association of LEDGF to chromosomes ^17,20,23,24^. However, we did find that the PWWP domain promoted binding within transcribed sequences. In an independent test of the role H3K36me3 may play in the function of LEDGF we examined integration behavior in HEK293T cells lacking expression of SETD2, the sole histone methyltransferase responsible for H3K36me3 (Suppl. Fig. S9) ^47,48^. Here too, we did not see patterns that resembles LEDGF independent integration (Suppl. Fig. S6, compare SETD2 KO to LEDGF KO and SETD2 Control) and integration levels per gene were not significantly altered in the absence of H3K36me3 (Fig. 6F). The heatmap of the top 1,000 integration targets showed little change in cells with the SETD2 KO (Fig. 6G). We conducted RIG-RAG analysis and found that integration in cells lacking H3K36me3 showed no significant change in specificity (Suppl Table S1). And evaluation of integration in SPIN states showed no observable changes in integration behavior (Supp. Table S2). The metagene plot from cells lacking SETD2 exhibited the typical shape of integration across gene bodies but in this case, there was a modest yet reproducible shift towards the start of transcription (Fig. 6H). Such a shift upstream might be expected if H3K36me3 makes a partial contribution to the distribution of LEDGF towards the 3’ end of gene bodies which is where H3K36me3 is concentrated. Regardless, these data showed that H3K36me3 did not make a substantial contribution to integration specificity of genes.

## Discussion

LEDGF is a ubiquitously expressed nuclear protein thought to function as a transcription co-activator ^29,49–56^. Depending on threshold parameters, our genome-wide study of transcription found hundreds to thousands of genes changed expression levels in cells lacking LEDGF (Figs. 1D and 2I). Our observation that LEDGF predominantly associates with TSSs of actively transcribed genes and recruits RNA Pol II provides direct support for the model that it functions globally as a co-activator. Our ChIPseq maps of tagged LEDGF with pronounced TSS binding differ from recent results of native chromatin methods that report LEDGF positioned more evenly spread across transcribed and untranscribed sequences ^28,57^. This difference is possibly due to our use of fixation before cell lysis which can trap complexes that may be lost during extraction.

Although the distinct location of LEDGF at TSSs is expected for a transcriptional co-activator, this is surprising for a chromatin binding protein with a PWWP domain that associates with nucleosomes specifically containing histone H3K36me3, a modification concentrated in the 3’ region of transcribed sequences ^37^. That LEDGF lacking the PWWP domain retained strong TSS association argues that this binding is unrelated to H3K36me3. However, removal of the PWWP domain resulted in reduced enrichment within transcribed sequences suggesting that some association of LEDGF across genes is due to interaction with H3K36me3. The profile of LEDGF-dependent HIV-1 integration sites across transcribed sequences is clear evidence that a portion of LEDGF protein does associate with regions containing H3K36me3. Our integration data with cells lacking SETD2 demonstrated that the PWWP-H3k36me3 interaction played little if any role in selecting specific genes for integration. Nevertheless, the H3K36me3 modification played a measurable role in distributing integration across the body of genes. These data together with the ChIP-seq data are consistent with H3K36me3-PWWP interactions contributing to LEDGF enrichment in downstream transcribed sequences.

Fusion of the LEDGF IBD to readers of specific chromatin features such as CpG islands, H3K9me3, H3K27me3, and H3K4me3 redirects the position of integration to the chromatin specified by the reader ^58–61^. These results indicate that the association of IBD to chromatin is sufficient to tether IN and position integration. It is therefore surprising that the prominent binding of LEDGF to promoters results in little integration at TSSs. An explanation is suggested by the lack of TSS binding when the IBD is removed (3XFLAG-LEDGF-1-325). This lack of TSS binding suggests that the IBD is associated with another factor that recruits LEDGF to TSSs. As a result, IN is prevented from binding LEDGF and mediate integration at TSSs. This possibility is supported by the result that MLL1, a factor demonstrated to form a structured complex with the IBD, binds TSSs where LEDGF is positioned. Our finding that MLL1 contributes significantly to LEDGF binding at TSSs genome-wide, and that LEDGF reciprocates by supporting MLL1 binding broadly, provides firm support for the model that MLL1 associates with the IBD at TSSs and prevents integration by outcompeting IN (Fig. 7).

**Fig. 7.**
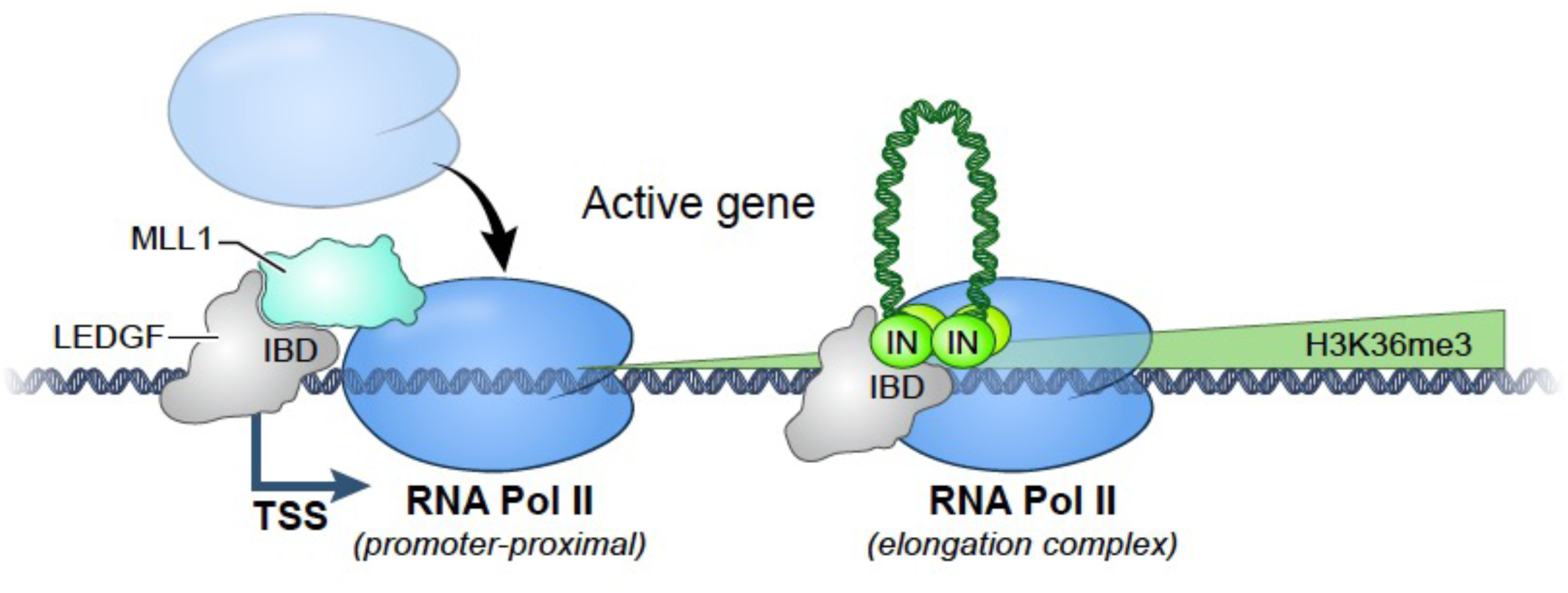
Model for the distribution of integration across actively transcribed genes. At TSSs MLL1 and LEDGF associate and recruit RNA Pol II. The IBD of LEDGF is bound by MLL1 blocking the association of IN. During elongation the IBD is available to interact with IN allowing integration (inverted U) to occur across transcribed sequences.

Biochemical evidence indicates that LEDGF promotes transcription elongation by functioning as a histone chaperone ^28^. In addition, elongation of RNA Pol II is regulated by a number of factors that interact with LEDGF ^21,22,62^. IWS1, SPT6, TFIIS, PPP1R10, and LEDGF form a quinary complex that contributes to RNA Pol II activity ^22^. We propose that LEDGF possesses dual functions. At promoters, LEDGF is a transcription co-activator where it recruits RNA Pol II to active promoters. During elongation, LEDGF leaves the promoter and in association with elongation factors it supports RNA Pol II activity across transcribed sequences. The transition of LEDGF from its association with TSSs to the elongation complex may be accompanied by transfer of the IBD from binding MLL1 to an association with the elongation factors. We speculate that integration occurs across transcribed sequences and not at TSSs because IN is unable to compete with MLL1 for IBD binding but succeeds when LEDGF participates in elongation (Fig. 7). Support for this proposal is the unique enrichment of MLL1 across HOX gene bodies, the strong co-accumulation of LEDGF at these regions, and the surprisingly low level of integration across HOX genes (Suppl. Fig. S8). Here too, MLL1 may prevent IN from associating with LEDGF.

MLL1 is the mammalian homolog of *Drosophila trithorax* and is a master regulator of HOX genes, which play a central role in embryogenesis and normal hematopoietic differentiation ^39,63–65^. Rearrangements of MLL1 occur in >70% of infant acute leukemia and in about 10% of acute myeloid leukemia (AML) cases in adults ^30–32^. The most common recombination events are fusions of the N-terminus of MLL1 with over 130 different partners producing oncogenic MLL1-fusion proteins (MLL1-FPs) ^66,67^. The MLL1-FPs create aberrant transcription by recruiting activating or elongation factors to MLL1 target genes and drive overexpression of *HOX* genes, a key feature thought to drive MLL leukemia pathogenesis ^31,66,68–71^. Importantly, the role played by MLL1-FPs in leukemic transformation depends on LEDGF ^41,42,56,72^. The MLL1-FPs retain the N-terminus of MLL1, which forms a triple complex with MENIN and LEDGF ^38,56,73–76^. The predominant binding of Menin to TSSs ^77–79^, the strong correlation of MLL1 and LEDGF at TSSs (Figs. 4 and 5), and our finding that LEDGF supports MLL1 association with TSSs genome-wide, provides a mechanistic explanation for the contribution of LEDGF to the oncogenic activity of MLL1-FPs.

## Methods

### Cells and viruses

HEK293T cells were propagated in Dulbecco’s modified Eagle’s media supplemented to contain fetal bovine serum (10%) and penicillin-streptomycin (1%; 100 U/ml) (DMEM). Cells were grown at 37 °C in humified tissue culture cell incubators.

HEK293T cells were modified with CRIPSR to prevent expression of both the p52 and p75 isoforms of LEDGF. Cas9 with two sgRNAs targeting exon 2 of PSIP1 generated a clone with one allele encoding a frameshift after amino acid 30 adding seven additional amino acids and a second allele with a stop codon at amino acid 24. RNA-seq data from this LEDGF KO cell line revealed nonsense mediated decay allowed only trace levels of expression at any position in the *PSIP1* gene. A clone of HEK293T with homozygous alleles of *PSIP1* encoding C-terminal 3XFLAG tags were produced with CRISPR and a donor sequence carrying only a silent mutation that removed the PAM sequence.

We expressed N-terminal tagged LEDGF from Sleeping Beauty vectors introduced into the HEK293T cells with the LEDGF KO. A Sleeping Beauty vector, a gift from Todd McFarlan, containing 5’ and 3’ terminal inverted repeats, puro and a PGK promoter to express coding sequence with an N-terminal 3XFLAG tag. At 5’ NcoI and 3’ NheI sites we introduced synthesized coding sequences for wild type LEDGF, LEDGF lacking amino acids 1-93 (deletes PWWP) and LEDGF-1-325 (deletes IBD). We co-transfected these vectors with pCMV-SB100X that expresses the Sleeping Beauty transposase that stably integrated the LEDGF sequences.

To knock down expression of MLL1 we used a set of 3 SMARTvector Lentiviral Human KMT2A hCMV-TurboGFP shRNA vectors from Horizon Discovery Biosciences Limited following the instructions provided with the kit. The SETD2 KO cells were the kind gift of Dr. W. Kimryn Rathmell and were previously described ^48^.

A single-round reporter virus derivative of HIV-1 strain NL4-3 containing heterologous sequences from the U3 region of equine infectious anemia virus embedded within the HIV-1 U3 region was made by co-transfecting HEK293T cells with pNLX.Luc.R-U3-tag ^80^ and pCG-VSV-G ^81^ at the mass ratio of 6:1. After 2 days, 0.45 µm-filtered cell supernatants concentrated by ultracentrifugation at 53,000 g for 2 h at 4 °C were treated with TURBO DNase at 37 °C for 1 h. Virus yield was assessed by p24 ELISA (Advanced Biotechnology Laboratories). The embedded U3 tag affords the ability to score de novo integration sites in cells containing pre-existing HIV-1 sequences, such as the MLL1 knockdown cells.

### RNA-seq

RNA was extracted from HEK293T and LEDGF KO cells with RNeasy Plus Mini kit (Qiagen) following manufacturer’s instructions. Library preparation and sequencing was conducted by the NICHD Molecular Genomics Core. Approximately 1 µg of total RNA per sample was purified with PolyA extraction, and then purified mRNAs were used to make RNA-Seq libraries with specific barcodes using Illumina TruSeq Stranded mRNA Library Prep Kit. All the RNA-Seq libraries were pooled together and sequenced using illumina NovaSeq 6000 to generate approximately ∼40 million 2×100 paired-end reads for each sample. The raw data were demultiplexed and analyzed further.

Raw sequence reads were processed using the same methods described below for ChIPseq sequence reads with the following parameter differences: cutadapt parameters “-a AGATCGGAAGAGCACACGTCTGAACTCCAGTCA -A AGATCGGAAGAGCGTCGTGTAGGGAAAGAGTGT -q 20 –minimum-length=25” for paired-end reads; PCR duplicates were flagged with picard MarkDuplicates with parameter “VALIDATION_STRINGENCY=LENIENT”; trimmed reads were mapped to the genome with hisat2 v 2.2.1 with the parameters “--no-unal -S”; Bigwig files were generated with deeptools bamCoverage with parameters “--minMappingQuality 20 --ignoreDuplicates --smoothlength”. Gene expression was then quantified using the subread featureCounts v2.0.1 with default parameters with the uniquely aligned reads and the reference annotation v19_GRCh37.87_merged_transcripts. The Gencode human reference genome release_19 GRCh37.p13 was used, along with the corresponding Gencode v19 GRCh37.87 GTF annotation. The annotation was further processed by filtering for transcript entries, and merging transcripts per gene with the R GenomicRanges reduce function, which we refer as v19_GRCh37.87_merged_transcripts.

Differential expression analysis was performed with DESeq2 v1.30.0.

### DEX-seq and DRIM-seq

From this analysis we wished to identify the contribution of LEDGF to splice choices. We identified Differential Usage Transcripts (DUTs) when we compare RNA from LEDGF KO to RNA of WT HEK293T cells. DEX-seq evaluates RNA-seq data from two samples to identify genes with isoforms that are differentially expressed ^82^. DRIM-seq seeks to identify differences in exon usage ^83^. We applied a pipeline as described which uses transcript isoform quantifications from Salmon as identical input for both DRIM-Seq and DEX-Seq analysis ^84^. Default parameters were used with a p value threshold of 0.05. The versions of these pipelines used for our analyses were DEX-seq (version 1.40.0) and DRIM-seq (version 1.22.0).

### ChIP-seq

Antibodies for ChIPSeq: FLAG (Sigma; Cat# F1804), RNA Pol II (8WG16 BioLegend; Cat# 664912), H3K4me3 (C42D8 Rabbit mAb #9751 cell signaling), H3K36me3 (Invitrogen; MA5-24687), and MLL1 (Rabbit anti-MLL1 antibody Bethyl; Cat#A300-374A).

The ChIP-seq protocol is divided into individual days of the procedure.

### Day 1: Formaldehyde cross-linking of samples and Chromatin isolation

(Chromatin for each experiment comes from 1-2 x 150mm dish fully confluent).

A. Cross-link directly in the dish with adherent cells adding formaldehyde slowly to the culture dish with gentle mixing, final concentration 2%. (Use 550μl of 36% stock per 10mL of media). Incubate room temp with gentle shaking for 10 min.
B. Add Glycine slowly to a final concentration of (stock is 2.5M Invitrogen 15527013) to stop the reaction. Incubate RT with gentle shaking for 10 min.
C. Wash twice with 20 ml cold PBS and remove PBS.
D. Incubate the plate on ice and add while gently stirring add 5 ml of cell lysis buffer dropwise. Incubate on ice for 2-3 min.
E. Scrape the cells and collect with lysis buffer in 15 ml screw cap conical tube.
F. Incubate the tube on ice for 15min.
G. centrifuge at 1200 rpm in JS5.3 rotor for 10 min at 4C.
H. Remove the supernatant and resuspend the pellet in 600μl of Nuclear Lysis Buffer and incubate on ice for 10 min, then freeze in liquid nitrogen and store in the -80°.

### Day 2: Chromatin sonication (Day 1 and Day 2 can be done on the same day if you don’t freeze the nuclear lysis)

Defrost the samples on ice from the -80°. If there are precipitates dissolve by gentle warming (this affects the efficiency of the sonication).

A. One hour before starting cool down the diagenode sonicator bath to 4C before sonication.
B. Once all the salt and precipitates are dissolved move the samples into a diagenode 1.5 ml glass tube and close the lid.
C. Set the machine for 30 cycles of 30 sec ON-30 sec OFF.
D. Move the samples to regular 1.5 ml eppendorf tube and centrifuge at max speed at 4° C for 10 min.
E. Collect the supernatant to a clean 2mL eppendorf tube, usually about 300 to 400 microliters.

### Day 3: Immunoprecipitation

A. Beads preparation

a. Mix 500μl each of Dyanabeads protein A (Thermo Fisher catalog no. 10002D) and dyanabeads protein G (Thermo Fisher catalog no. 10004D)
b. Add 100μl of the mixed beads in a 2ml tube for each of samples you will IP.
c. Set on magnet for 2 min.
d. Wash with 1 ml filter sterilized PBS-BSA (5mg/ml) X2
e. Resuspend beads in 300μl PBS-BSA solution.
f. Add 3-5μl antibody to each tube of beads.
g. Incubate at 4C on a rotating platform for min 6 hours with a max of overnight.
B. Chromatin preparation (precleared chromatin)

a. Prepare a mixture of Dyanabeads protein A and Dyanabeads protein G (20 μl for each of the samples you have) and with magnetic stand wash once with 1 ml of cold PBS-BSA (5 mg/ml) and once with 500 μl of ice-cold IP Dilution buffer (IPDB).
b. Dilute chromatin 10-fold with ice-cold IPDB. Replicas are made from post-sonication supernatants, usually 100 microliters plus 0.9 ml IPDB into one 2 ml tube per replicate.
c. Mix the 20 μl of beads with diluted chromatin.
d. Incubate on a rotating platform at 4°C for 2 hours.
e. After 2 hours, separate the diluted chromatin from the beads with a magnetic stand and transfer supernatant to a fresh 2 ml Eppendorf tube. This is known as precleared chromatin.
C. Wash with magnetic stand each tube of antibody bound beads from A above one time with 500μl of IPDB and discard supernatant.
D. Add the 1 ml precleared chromatin to the 2 ml tube containing antibody bound beads.
E. Keep on a rotating platform at 4°C overnight.

### Day4: Capture antibody/chromatin complex and reverse cross-links

A. Put the beads on a magnet and discard the supernatant.
B. Wash 10 times with the following cold buffers with 1ml each, mix briefly and keep rotating for 5 min each in a cold room.

2 x COLD IP wash Buffer I
3 x COLD IP High Salt Wash Buffer II
3 x COLD IP Wash Buffer III
2 x COLD TE Buffer
C. Elute antibody/chromatin complexes by adding 100 μl freshly made IP elution buffer. Mix well and incubate for 30 minutes at 65°C, be sure there is no condensation on lid. Collect the supernatant into a new 2 ml tube.
D. Add 100 μl freshly made IP elution buffer one more time onto the magnetic bead. Mix well and incubate for 30 minutes at 65°C. Combine the supernatants into the 2 ml tube above. Total volume is 200ul for each IP.
E. Mix gently and add 5 ul of 20mg/ml proteinase K (NEB) solution to each tube and incubate for 3 hr at 45°C being careful there is no condensation.
F. Add 24 μl of 5M NaCl (to give 0.6 M final concentration) to each tube.
G. Incubate all samples in a 65°C overnight in a heated lid PCR machine to reverse formaldehyde cross-links.

### Day5: ChIP DNA purification

A. Allow samples to cool down, add 8 ul of RNAse A (NEB, 10 mg/ml), and incubate 1 h at 37 C being careful there is no condensation on lid.
B. Use a chip purification kit (Zymo Research, Catalog no. D5205) to purify ChIP DNA.
C. Check the concentration by qubet.

### ChIP Buffers

**1. Cell Lysis Buffer (CLB) (Make same day as experiment)**

10 mM Tris-HCI pH 8.0
10 mM NaCI
0.2% NP40
1X protease inhibitors
1mM PMSF
**2. Nuclear Lysis Buffer (NLB): (Make same day as experiment)**

50 mM Tris pH 8.0
10mM EDTA pH 8.0
1% SDS
1X protease inhibitors
1mM PMSF
**3. IP Dilution Buffer (IPDB) (Make same day as experiment)**

20 mM Tris pH 8.0
2 mM EDTA
150 mM NaCI
1 % Triton X - 100
0.01% SDS
1X protease inhibitors
ImM PMSF
**4. IP Wash Buffer I (Filter sterilized Stored in fridge)**

20 mM Tris pH 8
2 mM EDTA
50 mM NaCI
1% Triton X - 100
0.1% SDS
**5. IP High Salt Wash Buffer II (Filter sterilized Stored in fridge)**

20 mM Tris pH 8
2 mM EDTA
500 mM NaCI
1% Triton X - 100
0.1% SDS
**6. IP Wash Buffer III (Filter sterilized stored in fridge)**

10 mM Tris pH 8
1 mM EDTA
250 mM LiCI
1% NP40
1% Deoxycholate
7. IX TE
10 mM Tris pH 8
1 mM EDTA
**8. Elution Buffer (made same day as experiment)**

100mM NaHCO3
1% SDS

Library production and sequencing was performed by the NIH Intramural Sequencing Center (NISC). ChIP-Seq libraries were constructed from 40 ng of ChIP DNA using Ovation Ultralow System V2 1-96 (Tecan) with 15 cycles of PCR amplification. The final libraries were twice purified using Ampure XP PCR Purification Beads (Agencourt). The pooled libraries were sequenced on a NovaSeq 6000 (Illumina) using version 1.5 chemistry to achieve approximately 80 million 151-base read pairs. The data was processed using RTA version 3.4.4.

Raw sequence reads were processed through lcdb-wf v1.6rc (https://github.com/lcdb/lcdb-wf). Briefly, raw sequence reads were trimmed with cutadapt v3.4 to remove adapters and do quality trimming with parameters “-u 3 -a AGATCGGAAGAGCACACGTCTGAACTCCAGTCA -A AGATCGGAAGAGCGTCGTGTAGGGAAAGAGTGT -q 20 –minimum-length=25” for paired-end reads or “-a AGATCGGAAGAGCACACGTCTGAACTCCAGTCA -q 20 – minimum-length=25” for single read. fastqc v0.11.9 with default parameters and multiqc v1.10.1 were used to estimate the sequencing library quality. The presence of common sequencing contaminants was evaluated with fastq_screen v0.14.0 with parameters “–subset 100000 –aligner bowtie2.” Trimmed reads were mapped to the human reference genome Gencode v19 GRCh37.p13) using bowtie2 v2.4.2 with parameter “--no-unal”, followed by samtools v1.12 functions view with parameter “-Sb” and function sort to sort the BAM files. Multimapping reads were filtered using samtools view with parameter “-b -q 20”. PCR duplicates were removed with picard v2.25.2 MarkDuplicates with parameters “REMOVE_DUPLICATES=true VALIDATION_STRINGENCY=LENIENT”. Bigwig files were generated with deeptools v3.5.1 bamCoverage with “--binSize 1 --minMappingQuality 20 --ignoreDuplicates --normalizeUsing CPM --extendReads 300” to visualize the signal in genome browsers. Significant ChIPseq signal peaks were computed with MACS2 v2.2.7.1 callpeak with parameters “-f BAMPE -B” or “-f BAM -B” for paired-end or single read respectively. The “-B” parameter was used to output the bedGraph files required to return the fold enrichment (FE) of IP over input BigWig files with MACS2 bdgcmp with parameter “-m FE”.

The list of genes containing LEDGF-3XFLAG peaks was generated by converting in R the MACS2 peaks to a GenomicRanges v1.50.0 Granges object and annotating the peaks with a fold-enrichment > 2.5 with the ChIPpeakAnno v3.32.0 function annoPeaks with the parameters “bindingType=’fullRange’, bindingRegion=c(0,1)”.

Random binding would be expected to have a genome-wide fold-enrichment of 1; random datasets were generated by assigning a value of 1 to all regions mapped in the corresponding sample. The random controls were then processed identically to the experimental signals.

Matrices of FE in bins across the regions 8kb upstream of genes to 8kb downstream were calculated with deeptools v3.5.1 computeMatrix scale-regions with the same parameters than the HIV insertion matrices besides “--averageTypeBins max”, which is recommended for ChIPseq. The resulting matrices were summarized and plotted in R with the same functions than the HIV insertion metagene plots. Metagene plots were done for either all genes, subsetted to the genes containing LEDGF-3XFLAG, or subsetted to the differentially expressed genes (FDR < 0.1, abs(LFC) > 0 or > 0.7 as indicated).

#### Metagene representations

Matrices of fold enrichment in bins across the regions 8kb upstream of genes to 8kb downstream were calculated with deeptools v3.5.1 computeMatrix scale-regions with the same parameters used with the HIV insertion matrices besides “--averageTypeBins max”, which is recommended for ChIPseq. The resulting matrices were summarized and plotted in R: averages and standard errors (SE) across genes of the sums of insertions per gene and per bin were calculated with the r-base v4.0.2 function colMeans. Metagene graphs were plotted with the R ggplot2 v3.3.3 functions geom_line and geom_ribbon for the average insertion sum per bin and SE respectively. Metagene plots were done for either all genes, subsetted to the genes containing LEDGF-3XFLAG, or subsetted to the differentially expressed genes (FDR < 0.1, abs(LFC) > 0 or > 0.7 as indicated).

#### Integration sequencing

HEK293T cells were infected with HIV-1_NLX.Luc.R-U3-tag_ at 0.5 pg p24/cell. After 5 h, virus was removed, and cell cultures were replenished with fresh DMEM. Infections proceeded for 5 d. Integration site sequencing libraries were prepared essentially as previously described in Serrao et al. ^85^. Briefly, genomic DNA (5 μg) digested overnight with a cocktail of enzymes (AvrII, NheI-HF, SpeI-HF, and BamH1-HF; 100 units each, was purified using a PCR purification kit. The purified DNA was ligated with freshly annealed asymmetric linkers overnight as four parallel reactions and purified using a PCR purification kit. The ligated samples were subjected to two rounds of ligation mediated (LM)-PCR. After PCR purification, libraries were quantified by DNA fluorimetry (Qubit) and pooled at 10nM. PhiX174 control DNA was spiked-in to the pooled libraries at 30% and the mixture was loaded into a P2 300 cycle cartridge at 650 pM concentration and sequenced on an Illumina NextSeq2000.

Raw fastq files were demultiplexed using Sabre tool. Post demultiplexing, the files were trimmed, aligned to hg19 genome and bed files were generated as described ^85^. Integration into genes and SPADs was scored as within these genomic coordinates. For TSSs, CpG islands, LADs, H3K4me3, and H3K36me3, sites were mapped within +/-2.5 kb windows (5 kb surrounding these coordinates). Gene Density (GD) is the number of genes in +/-500 kb (1 Mb window). Further analysis was performed at the NICHD Bioinformatics and Scientific Programming Core. The Gencode human reference genome release_19 GRCh37.p13 was used, along with the corresponding Gencode v19 GRCh37.87 GTF annotation. The annotation was further processed by filtering for transcript entries, and merging transcripts per gene with the R GenomicRanges reduce function, which we refer as v19_GRCh37.87_merged_transcripts. HIV insertions were listed in BED-formatted files. Bigwig format is required to map the insertions to the genes with deeptools v3.5.1 computematrix. BED files were first converted to bedGraph by reformatting using pandas dataframes. Eventual overlapping insertions were merged with bedtools v2.30.0 merge with parameters “-c 4 -o sum”, then bigwig files were created with ucsc bedGraphToBigWig v377. Matrices of insertions mapped to the regions 8kb upstream of genes to 8kb downstream were calculated with deeptools v3.5.1 computeMatrix with parameters “scale-regions --binsize 200 --averageTypeBins sum --beforeRegionStartLength 8000 -- afterRegionStartLength 8000 --regionBodyLength 8000 --missingDataAsZero”. A random control insertion profile was generated and processed identically to the experimental HIV insertion profiles. The random integration control (RIC) was generated in silico based on the genomic DNA fragmentation technique ^86^.

The resulting matrices were summarized and plotted in R: the sums of insertions per gene and per bin were averaged across the genes with the r-base v4.0.2 function colMeans and the standard errors (SE) calculated. The metagene plots were generated with the R ggplot2 v3.3.3 functions geom_line and geom_ribbon for the average insertion sum per bin and SE respectively. Significant differences in proportion of HIV insertions along the genes were tested by dividing each ORF in 15 equal-length bins. Bins were generated with pybedtools v0.8.2 window_maker. Negative strand ORF bins’ order was inverted to have bin 1 at the 5’ end of the ORF. The 8kb upstream and downstream regions were processed similarly. For each gene and each bin, the percent of total insertions were quantified by counting the number of hits of pybedtools intersect with parameter “u=True”, and normalizing by total number of insertions. Biological replicates were merged to have a count matrix genes x conditions. T-tests and FDR p-value adjustments were performed in R with the functions t.test and p.adjust method=”BH”, first between biological replicates to confirm they are not statistically different and can be merged, then between condition to test the difference in proportion between SETD2_WT and SETD2_KO.

## Declarations

- Availability of data and materials

All data generated or analyzed during this study are included in this published article [and its supplementary information files].
- Declaration of interests

The authors declare that they have no competing interests.
- Funding

- Funding: This research was supported by the Intramural Research Program of the National Institutes of Health (NIH) from the *Eunice Kennedy Shriver* National Institute of Child Health and Human Development and the NIH Office of AIDS Research (HLL) and extramural NIH grant R37AI039394 (ANE).
- Authors’ contributions

Conceptualization: RP and HLL. Experimentation: RP, RR, PKS, AA, PS, and AR. Bioinformatic analysis: CE, PKS, HZ, RD, and GJB. Visualization: RP, CE, RR, RKS, and HLL. Supervision: SH, ANE and HLL. Writing original draft: HLL. Writing and editing: ANE and HLL.

- Supplemental information titles and legends

## Acknowledgements

This research was supported by the Intramural Research Program of the National Institutes of Health (NIH) from the *Eunice Kennedy Shriver* National Institute of Child Health and Human Development.

## References

1. Bedwell, G.J., and Engelman, A.N. (2021). Factors that mold the nuclear landscape of HIV-1 integration. Nucleic Acids Res 49, 621–635. 10.1093/nar/gkaa1207.

2. Coffin, J.M., and Hughes, S.H. (2021). Clonal Expansion of Infected CD4+ T Cells in People Living with HIV. Viruses 13. 10.3390/v13102078.

3. Maldarelli, F., Wu, X., Su, L., Simonetti, F.R., Shao, W., Hill, S., Spindler, J., Ferris, A.L., Mellors, J.W., Kearney, M.F., et al. (2014). HIV latency. Specific HIV integration sites are linked to clonal expansion and persistence of infected cells. Science. 345, 179–183. 10.1126/science.1254194.

4. Achuthan, V., Perreira, J.M., Sowd, G.A., Puray-Chavez, M., McDougall, W.M., Paulucci-Holthauzen, A., Wu, X., Fadel, H.J., Poeschla, E.M., Multani, A.S., et al. (2018). Capsid-CPSF6 Interaction Licenses Nuclear HIV-1 Trafficking to Sites of Viral DNA Integration. Cell Host Microbe 24, 392–404 e398. 10.1016/j.chom.2018.08.002.

5. Francis, A.C., Marin, M., Singh, P.K., Achuthan, V., Prellberg, M.J., Palermino-Rowland, K., Lan, S., Tedbury, P.R., Sarafianos, S.G., Engelman, A.N., and Melikyan, G.B. (2020). HIV-1 replication complexes accumulate in nuclear speckles and integrate into speckle-associated genomic domains. Nat Commun 11, 3505. 10.1038/s41467-020-17256-8.

6. Engelman, A.N. (2021). HIV Capsid and Integration Targeting. Viruses 13. 10.3390/v13010125.

7. Jang, S., and Engelman, A.N. (2023). Capsid-host interactions for HIV-1 ingress. Microbiol Mol Biol Rev, e0004822. 10.1128/mmbr.00048-22.

8. Schroder, A.R., Shinn, P., Chen, H., Berry, C., Ecker, J.R., and Bushman, F. (2002). HIV-1 integration in the human genome favors active genes and local hotspots. Cell 110, 521–529.

9. Llano, M., Vanegas, M., Fregoso, O., Saenz, D., Chung, S., Peretz, M., and Poeschla, E.M. (2004). LEDGF/p75 determines cellular trafficking of diverse lentiviral but not murine oncoretroviral integrase proteins and is a component of functional lentiviral preintegration complexes. Journal of Virology 78, 9524–9537.

10. Ciuffi, A., Llano, M., Poeschla, E., Hoffmann, C., Leipzig, J., Shinn, P., Ecker, J.R., and Bushman, F. (2005). A role for LEDGF/p75 in targeting HIV DNA integration. Nat Med. 11, 1287–1289. Epub 2005 Nov 1227.

11. Wang, G.P., Ciuffi, A., Leipzig, J., Berry, C.C., and Bushman, F.D. (2007). HIV integration site selection: analysis by massively parallel pyrosequencing reveals association with epigenetic modifications. Genome Res 17, 1186–1194.

12. Shun, M.C., Raghavendra, N.K., Vandegraaff, N., Daigle, J.E., Hughes, S., Kellam, P., Cherepanov, P., and Engelman, A. (2007). LEDGF/p75 functions downstream from preintegration complex formation to effect gene-specific HIV-1 integration. Genes Dev 21, 1767–1778. 21/14/1767 [pii] 10.1101/gad.1565107.

13. Sowd, G.A., Serrao, E., Wang, H., Wang, W., Fadel, H.J., Poeschla, E.M., and Engelman, A.N. (2016). A critical role for alternative polyadenylation factor CPSF6 in targeting HIV-1 integration to transcriptionally active chromatin. Proc Natl Acad Sci U S A 113, E1054–1063. 10.1073/pnas.1524213113.

14. Maertens, G.N., Engelman, A.N., and Cherepanov, P. (2022). Structure and function of retroviral integrase. Nat Rev Microbiol 20, 20–34. 10.1038/s41579-021-00586-9.

15. Bedwell, G.J., Jang, S., Li, W., Singh, P.K., and Engelman, A.N. (2021). rigrag: high-resolution mapping of genic targeting preferences during HIV-1 integration in vitro and in vivo. Nucleic Acids Res 49, 7330–7346. 10.1093/nar/gkab514.

16. Cherepanov, P., Devroe, E., Silver, P.A., and Engelman, A. (2004). Identification of an evolutionarily conserved domain in human lens epithelium-derived growth factor/transcriptional co-activator p75 (LEDGF/p75) that binds HIV-1 integrase. J Biol Chem 279, 48883–48892. 10.1074/jbc.M406307200 M406307200 [pii].

17. Llano, M., Vanegas, M., Hutchins, N., Thompson, D., Delgado, S., and Poeschla, E.M. (2006). Identification and characterization of the chromatin-binding domains of the HIV-1 integrase interactor LEDGF/p75. J Mol Biol 360, 760–773. 10.1016/j.jmb.2006.04.073.

18. Turlure, F., Maertens, G., Rahman, S., Cherepanov, P., and Engelman, A. (2006). A tripartite DNA-binding element, comprised of the nuclear localization signal and two AT-hook motifs, mediates the association of LEDGF/p75 with chromatin in vivo. Nucleic Acids Res 34, 1653–1665. 34/5/1663 [pii] 10.1093/nar/gkl052.

19. Pradeepa, M.M., Sutherland, H.G., Ule, J., Grimes, G.R., and Bickmore, W.A. (2012). Psip1/Ledgf p52 binds methylated histone H3K36 and splicing factors and contributes to the regulation of alternative splicing. PLoS Genet 8, e1002717. 10.1371/journal.pgen.1002717.

20. Eidahl, J.O., Crowe, B.L., North, J.A., McKee, C.J., Shkriabai, N., Feng, L., Plumb, M., Graham, R.L., Gorelick, R.J., Hess, S., et al. (2013). Structural basis for high-affinity binding of LEDGF PWWP to mononucleosomes. Nucleic Acids Res. 41, 3924–3936. 10.1093/nar/gkt074.

21. Tesina, P., Cermakova, K., Horejsi, M., Prochazkova, K., Fabry, M., Sharma, S., Christ, F., Demeulemeester, J., Debyser, Z., De Rijck, J., et al. (2015). Multiple cellular proteins interact with LEDGF/p75 through a conserved unstructured consensus motif. Nat Commun 6, 7968. 10.1038/ncomms8968.

22. Cermakova, K., Demeulemeester, J., Lux, V., Nedomova, M., Goldman, S.R., Smith, E.A., Srb, P., Hexnerova, R., Fabry, M., Madlikova, M., et al. (2021). A ubiquitous disordered protein interaction module orchestrates transcription elongation. Science 374, 1113–1121. 10.1126/science.abe2913.

23. van Nuland, R., van Schaik, F.M., Simonis, M., van Heesch, S., Cuppen, E., Boelens, R., Timmers, H.M., and van Ingen, H. (2013). Nucleosomal DNA binding drives the recognition of H3K36-methylated nucleosomes by the PSIP1-PWWP domain. Epigenetics Chromatin 6, 12. 10.1186/1756-8935-6-12.

24. Wang, H., Farnung, L., Dienemann, C., and Cramer, P. (2020). Structure of H3K36-methylated nucleosome-PWWP complex reveals multivalent cross-gyre binding. Nat Struct Mol Biol 27, 8–13. 10.1038/s41594-019-0345-4.

25. Sapp, N., Burge, N., Cox, K., Prakash, P., Balasubramaniam, M., Thapa, S., Christensen, D., Li, M., Linderberger, J., Kvaratskhelia, M., et al. (2022). HIV-1 Preintegration Complex Preferentially Integrates the Viral DNA into Nucleosomes Containing Trimethylated Histone 3-Lysine 36 Modification and Flanking Linker DNA. J Virol 96, e0101122. 10.1128/jvi.01011-22.

26. Wang, H., Jurado, K.A., Wu, X., Shun, M.C., Li, X., Ferris, A.L., Smith, S.J., Patel, P.A., Fuchs, J.R., Cherepanov, P., et al. (2012). HRP2 determines the efficiency and specificity of HIV-1 integration in LEDGF/p75 knockout cells but does not contribute to the antiviral activity of a potent LEDGF/p75-binding site integrase inhibitor. Nucleic Acids Res 40, 11518–11530. 10.1093/nar/gks913.

27. Singh, P.K., Plumb, M.R., Ferris, A.L., Iben, J.R., Wu, X., Fadel, H.J., Luke, B.T., Esnault, C., Poeschla, E.M., Hughes, S.H., et al. (2015). LEDGF/p75 interacts with mRNA splicing factors and targets HIV-1 integration to highly spliced genes. Genes Dev 29, 2287–2297. 10.1101/gad.267609.115.

28. LeRoy, G., Oksuz, O., Descostes, N., Aoi, Y., Ganai, R.A., Kara, H.O., Yu, J.R., Lee, C.H., Stafford, J., Shilatifard, A., and Reinberg, D. (2019). LEDGF and HDGF2 relieve the nucleosome-induced barrier to transcription in differentiated cells. Sci Adv 5, eaay3068. 10.1126/sciadv.aay3068.

29. Ge, H., Si, Y., and Roeder, R.G. (1998). Isolation of cDNAs encoding novel transcription coactivators p52 and p75 reveals an alternate regulatory mechanism of transcriptional activation. EMBO J 17, 6723–6729. 10.1093/emboj/17.22.6723.

30. Chen, C.S., Sorensen, P.H., Domer, P.H., Reaman, G.H., Korsmeyer, S.J., Heerema, N.A., Hammond, G.D., and Kersey, J.H. (1993). Molecular rearrangements on chromosome 11q23 predominate in infant acute lymphoblastic leukemia and are associated with specific biologic variables and poor outcome. Blood 81, 2386–2393.

31. Mohan, M., Lin, C., Guest, E., and Shilatifard, A. (2010). Licensed to elongate: a molecular mechanism for MLL-based leukaemogenesis. Nat Rev Cancer 10, 721–728. 10.1038/nrc2915.

32. Issa, G.C., Ravandi, F., DiNardo, C.D., Jabbour, E., Kantarjian, H.M., and Andreeff, M. (2021). Therapeutic implications of menin inhibition in acute leukemias. Leukemia 35, 2482–2495. 10.1038/s41375-021-01309-y.

33. Santos-Rosa, H., Schneider, R., Bannister, A.J., Sherriff, J., Bernstein, B.E., Emre, N.C., Schreiber, S.L., Mellor, J., and Kouzarides, T. (2002). Active genes are tri-methylated at K4 of histone H3. Nature 419, 407–411. 10.1038/nature01080.

34. Bernstein, B.E., Kamal, M., Lindblad-Toh, K., Bekiranov, S., Bailey, D.K., Huebert, D.J., McMahon, S., Karlsson, E.K., Kulbokas, E.J., 3rd, Gingeras, T.R., et al. (2005). Genomic maps and comparative analysis of histone modifications in human and mouse. Cell 120, 169–181. 10.1016/j.cell.2005.01.001.

35. Piunti, A., and Shilatifard, A. (2016). Epigenetic balance of gene expression by Polycomb and COMPASS families. Science 352, aad9780. 10.1126/science.aad9780.

36. Wang, H., Fan, Z., Shliaha, P.V., Miele, M., Hendrickson, R.C., Jiang, X., and Helin, K. (2023). H3K4me3 regulates RNA polymerase II promoter-proximal pause-release. Nature 615, 339–348. 10.1038/s41586-023-05780-8.

37. Wagner, E.J., and Carpenter, P.B. (2012). Understanding the language of Lys36 methylation at histone H3. Nat Rev Mol Cell Biol 13, 115–126. 10.1038/nrm3274.

38. Huang, J., Gurung, B., Wan, B., Matkar, S., Veniaminova, N.A., Wan, K., Merchant, J.L., Hua, X., and Lei, M. (2012). The same pocket in menin binds both MLL and JUND but has opposite effects on transcription. Nature 482, 542–546. 10.1038/nature10806.

39. Yu, B.D., Hess, J.L., Horning, S.E., Brown, G.A., and Korsmeyer, S.J. (1995). Altered Hox expression and segmental identity in Mll-mutant mice. Nature 378, 505–508. 10.1038/378505a0.

40. Yokoyama, A., Wang, Z., Wysocka, J., Sanyal, M., Aufiero, D.J., Kitabayashi, I., Herr, W., and Cleary, M.L. (2004). Leukemia proto-oncoprotein MLL forms a SET1-like histone methyltransferase complex with menin to regulate Hox gene expression. Mol Cell Biol 24, 5639–5649. 10.1128/MCB.24.13.5639-5649.2004.

41. Pradeepa, M.M., Grimes, G.R., Taylor, G.C., Sutherland, H.G., and Bickmore, W.A. (2014). Psip1/Ledgf p75 restrains Hox gene expression by recruiting both trithorax and polycomb group proteins. Nucleic Acids Res 42, 9021–9032. 10.1093/nar/gku647.

42. El Ashkar, S., Schwaller, J., Pieters, T., Goossens, S., Demeulemeester, J., Christ, F., Van Belle, S., Juge, S., Boeckx, N., Engelman, A., et al. (2018). LEDGF/p75 is dispensable for hematopoiesis but essential for MLL-rearranged leukemogenesis. Blood 131, 95–107. 10.1182/blood-2017-05-786962.

43. Shilatifard, A. (2012). The COMPASS family of histone H3K4 methylases: mechanisms of regulation in development and disease pathogenesis. Annu Rev Biochem 81, 65–95. 10.1146/annurev-biochem-051710-134100.

44. Rao, R.C., and Dou, Y. (2015). Hijacked in cancer: the KMT2 (MLL) family of methyltransferases. Nat Rev Cancer 15, 334–346. 10.1038/nrc3929.

45. Llano, M., Saenz, D.T., Meehan, A., Wongthida, P., Peretz, M., Walker, W.H., Teo, W., and Poeschla, E.M. (2006). An essential role for LEDGF/p75 in HIV integration. Science 314, 461–464. 1132319 [pii] 10.1126/science.1132319.

46. Wang, Y., Zhang, Y., Zhang, R., van Schaik, T., Zhang, L., Sasaki, T., Peric-Hupkes, D., Chen, Y., Gilbert, D.M., van Steensel, B., et al. (2021). SPIN reveals genome-wide landscape of nuclear compartmentalization. Genome Biol 22, 36. 10.1186/s13059-020-02253-3.

47. Edmunds, J.W., Mahadevan, L.C., and Clayton, A.L. (2008). Dynamic histone H3 methylation during gene induction: HYPB/Setd2 mediates all H3K36 trimethylation. EMBO J 27, 406–420. 10.1038/sj.emboj.7601967.

48. Hacker, K.E., Fahey, C.C., Shinsky, S.A., Chiang, Y.J., DiFiore, J.V., Jha, D.K., Vo, A.H., Shavit, J.A., Davis, I.J., Strahl, B.D., and Rathmell, W.K. (2016). Structure/Function Analysis of Recurrent Mutations in SETD2 Protein Reveals a Critical and Conserved Role for a SET Domain Residue in Maintaining Protein Stability and Histone H3 Lys-36 Trimethylation. J Biol Chem 291, 21283–21295. 10.1074/jbc.M116.739375.

49. Fatma, N., Singh, D.P., Shinohara, T., and Chylack, L.T., Jr. (2001). Transcriptional regulation of the antioxidant protein 2 gene, a thiol-specific antioxidant, by lens epithelium-derived growth factor to protect cells from oxidative stress. J Biol Chem 276, 48899–48907. 10.1074/jbc.M100733200.

50. Kubo, E., Fatma, N., Sharma, P., Shinohara, T., Chylack, L.T., Jr., Akagi, Y., and Singh, D.P. (2002). Transactivation of involucrin, a marker of differentiation in keratinocytes, by lens epithelium-derived growth factor (LEDGF). J Mol Biol 320, 1053–1063. 10.1016/s0022-2836(02)00551-x.

51. Shinohara, T., Singh, D.P., and Fatma, N. (2002). LEDGF, a survival factor, activates stress-related genes. Prog Retin Eye Res 21, 341–358. 10.1016/s1350-9462(02)00007-1.

52. Sharma, P., Fatma, N., Kubo, E., Shinohara, T., Chylack, L.T., Jr., and Singh, D.P. (2003). Lens epithelium-derived growth factor relieves transforming growth factor-beta1-induced transcription repression of heat shock proteins in human lens epithelial cells. J Biol Chem 278, 20037–20046. 10.1074/jbc.M212016200.

53. Fatma, N., Kubo, E., Chylack, L.T., Jr., Shinohara, T., Akagi, Y., and Singh, D.P. (2004). LEDGF regulation of alcohol and aldehyde dehydrogenases in lens epithelial cells: stimulation of retinoic acid production and protection from ethanol toxicity. Am J Physiol Cell Physiol 287, C508–516. 10.1152/ajpcell.00076.2004.

54. Fatma, N., Kubo, E., Sharma, P., Beier, D.R., and Singh, D.P. (2005). Impaired homeostasis and phenotypic abnormalities in Prdx6-/-mice lens epithelial cells by reactive oxygen species: increased expression and activation of TGFbeta. Cell Death Differ 12, 734–750. 10.1038/sj.cdd.4401597.

55. Shin, J.H., Piao, C.S., Lim, C.M., and Lee, J.K. (2008). LEDGF binding to stress response element increases alphaB-crystallin expression in astrocytes with oxidative stress. Neurosci Lett 435, 131–136. 10.1016/j.neulet.2008.02.029.

56. Yokoyama, A., and Cleary, M.L. (2008). Menin critically links MLL proteins with LEDGF on cancer-associated target genes. Cancer Cell 14, 36–46. 10.1016/j.ccr.2008.05.003.

57. Jayakumar, S., Patel, M., Boulet, F., Aziz, H., Brooke, G.N., Tummala, H., and Pradeepa, M.M. (2024). PSIP1/LEDGF reduces R-loops at transcription sites to maintain genome integrity. Nat Commun 15, 361. 10.1038/s41467-023-44544-w.

58. Ferris, A.L., Wu, X., Hughes, C.M., Stewart, C., Smith, S.J., Milne, T.A., Wang, G.G., Shun, M.C., Allis, C.D., Engelman, A., and Hughes, S.H. (2010). Lens epithelium-derived growth factor fusion proteins redirect HIV-1 DNA integration. Proc Natl Acad Sci U S A 107, 3135–3140. 0914142107 [pii] 10.1073/pnas.0914142107.

59. Gijsbers, R., Ronen, K., Vets, S., Malani, N., De Rijck, J., McNeely, M., Bushman, F.D., and Debyser, Z. (2010). LEDGF hybrids efficiently retarget lentiviral integration into heterochromatin. Mol Ther 18, 552–560. 10.1038/mt.2010.36.

60. Silvers, R.M., Smith, J.A., Schowalter, M., Litwin, S., Liang, Z., Geary, K., and Daniel, R. (2010). Modification of integration site preferences of an HIV-1-based vector by expression of a novel synthetic protein. Hum Gene Ther 21, 337–349. 10.1089/hum.2009.134.

61. Tchasovnikarova, I.A., Marr, S.K., Damle, M., and Kingston, R.E. (2022). TRACE generates fluorescent human reporter cell lines to characterize epigenetic pathways. Mol Cell 82, 479–491 e477. 10.1016/j.molcel.2021.11.035.

62. Cermakova, K., Veverka, V., and Hodges, H.C. (2023). The TFIIS N-terminal domain (TND): a transcription assembly module at the interface of order and disorder. Biochem Soc Trans 51, 125–135. 10.1042/BST20220342.

63. Sauvageau, G., Lansdorp, P.M., Eaves, C.J., Hogge, D.E., Dragowska, W.H., Reid, D.S., Largman, C., Lawrence, H.J., and Humphries, R.K. (1994). Differential expression of homeobox genes in functionally distinct CD34+ subpopulations of human bone marrow cells. Proc Natl Acad Sci U S A 91, 12223–12227. 10.1073/pnas.91.25.12223.

64. Jude, C.D., Climer, L., Xu, D., Artinger, E., Fisher, J.K., and Ernst, P. (2007). Unique and independent roles for MLL in adult hematopoietic stem cells and progenitors. Cell Stem Cell 1, 324–337. 10.1016/j.stem.2007.05.019.

65. McMahon, K.A., Hiew, S.Y., Hadjur, S., Veiga-Fernandes, H., Menzel, U., Price, A.J., Kioussis, D., Williams, O., and Brady, H.J. (2007). Mll has a critical role in fetal and adult hematopoietic stem cell self-renewal. Cell Stem Cell 1, 338–345. 10.1016/j.stem.2007.07.002.

66. Hess, J.L. (2004). MLL: a histone methyltransferase disrupted in leukemia. Trends Mol Med 10, 500–507. 10.1016/j.molmed.2004.08.005.

67. Meyer, C., Burmeister, T., Groger, D., Tsaur, G., Fechina, L., Renneville, A., Sutton, R., Venn, N.C., Emerenciano, M., Pombo-de-Oliveira, M.S., et al. (2018). The MLL recombinome of acute leukemias in 2017. Leukemia 32, 273–284. 10.1038/leu.2017.213.

68. Daser, A., and Rabbitts, T.H. (2004). Extending the repertoire of the mixed-lineage leukemia gene MLL in leukemogenesis. Genes Dev 18, 965–974. 10.1101/gad.1195504.

69. Milne, T.A., Kim, J., Wang, G.G., Stadler, S.C., Basrur, V., Whitcomb, S.J., Wang, Z., Ruthenburg, A.J., Elenitoba-Johnson, K.S., Roeder, R.G., and Allis, C.D. (2010). Multiple interactions recruit MLL1 and MLL1 fusion proteins to the HOXA9 locus in leukemogenesis. Mol Cell 38, 853–863. 10.1016/j.molcel.2010.05.011.

70. Yokoyama, A., Lin, M., Naresh, A., Kitabayashi, I., and Cleary, M.L. (2010). A higher-order complex containing AF4 and ENL family proteins with P-TEFb facilitates oncogenic and physiologic MLL-dependent transcription. Cancer Cell 17, 198–212. 10.1016/j.ccr.2009.12.040.

71. Lin, C., Smith, E.R., Takahashi, H., Lai, K.C., Martin-Brown, S., Florens, L., Washburn, M.P., Conaway, J.W., Conaway, R.C., and Shilatifard, A. (2010). AFF4, a component of the ELL/P-TEFb elongation complex and a shared subunit of MLL chimeras, can link transcription elongation to leukemia. Mol Cell 37, 429–437. 10.1016/j.molcel.2010.01.026.

72. Mereau, H., De Rijck, J., Cermakova, K., Kutz, A., Juge, S., Demeulemeester, J., Gijsbers, R., Christ, F., Debyser, Z., and Schwaller, J. (2013). Impairing MLL-fusion gene-mediated transformation by dissecting critical interactions with the lens epithelium-derived growth factor (LEDGF/p75). Leukemia 27, 1245–1253. 10.1038/leu.2013.10.

73. Yokoyama, A., Somervaille, T.C., Smith, K.S., Rozenblatt-Rosen, O., Meyerson, M., and Cleary, M.L. (2005). The menin tumor suppressor protein is an essential oncogenic cofactor for MLL-associated leukemogenesis. Cell 123, 207–218. 10.1016/j.cell.2005.09.025.

74. Caslini, C., Yang, Z., El-Osta, M., Milne, T.A., Slany, R.K., and Hess, J.L. (2007). Interaction of MLL amino terminal sequences with menin is required for transformation. Cancer research 67, 7275–7283. 10.1158/0008-5472.CAN-06-2369.

75. Murai, M.J., Pollock, J., He, S., Miao, H., Purohit, T., Yokom, A., Hess, J.L., Muntean, A.G., Grembecka, J., and Cierpicki, T. (2014). The same site on the integrase-binding domain of lens epithelium-derived growth factor is a therapeutic target for MLL leukemia and HIV. Blood 124, 3730–3737. 10.1182/blood-2014-01-550079.

76. Cermakova, K., Tesina, P., Demeulemeester, J., El Ashkar, S., Mereau, H., Schwaller, J., Rezacova, P., Veverka, V., and De Rijck, J. (2014). Validation and structural characterization of the LEDGF/p75-MLL interface as a new target for the treatment of MLL-dependent leukemia. Cancer research 74, 5139–5151. 10.1158/0008-5472.CAN-13-3602.

77. Agarwal, S.K., Impey, S., McWeeney, S., Scacheri, P.C., Collins, F.S., Goodman, R.H., Spiegel, A.M., and Marx, S.J. (2007). Distribution of menin-occupied regions in chromatin specifies a broad role of menin in transcriptional regulation. Neoplasia 9, 101–107. 10.1593/neo.06706.

78. Cheng, J., Blum, R., Bowman, C., Hu, D., Shilatifard, A., Shen, S., and Dynlacht, B.D. (2014). A role for H3K4 monomethylation in gene repression and partitioning of chromatin readers. Mol Cell 53, 979–992. 10.1016/j.molcel.2014.02.032.

79. Scacheri, P.C., Davis, S., Odom, D.T., Crawford, G.E., Perkins, S., Halawi, M.J., Agarwal, S.K., Marx, S.J., Spiegel, A.M., Meltzer, P.S., and Collins, F.S. (2006). Genome-wide analysis of menin binding provides insights into MEN1 tumorigenesis. PLoS Genet 2, e51. 10.1371/journal.pgen.0020051.

80. Anderson-Daniels, J., Singh, P.K., Sowd, G.A., Li, W., Engelman, A.N., and Aiken, C. (2019). Dominant Negative MA-CA Fusion Protein Is Incorporated into HIV-1 Cores and Inhibits Nuclear Entry of Viral Preintegration Complexes. J Virol 93. 10.1128/JVI.01118-19.

81. Shun, M.C., Daigle, J.E., Vandegraaff, N., and Engelman, A. (2007). Wild-type levels of human immunodeficiency virus type 1 infectivity in the absence of cellular emerin protein. J Virol 81, 166–172. 10.1128/JVI.01953-06.

82. Anders, S., Reyes, A., and Huber, W. (2012). Detecting differential usage of exons from RNA-seq data. Genome Res 22, 2008–2017. 10.1101/gr.133744.111.

83. Nowicka, M., and Robinson, M.D. (2016). DRIMSeq: a Dirichlet-multinomial framework for multivariate count outcomes in genomics. F1000Res 5, 1356. 10.12688/f1000research.8900.2.

84. Love, M.I., Soneson, C., and Patro, R. (2018). Swimming downstream: statistical analysis of differential transcript usage following Salmon quantification. F1000Res 7, 952. 10.12688/f1000research.15398.3.

85. Serrao, E., Cherepanov, P., and Engelman, A.N. (2016). Amplification, Next-generation Sequencing, and Genomic DNA Mapping of Retroviral Integration Sites. J Vis Exp. 10.3791/53840.

86. Li, W., Singh, P.K., Sowd, G.A., Bedwell, G.J., Jang, S., Achuthan, V., Oleru, A.V., Wong, D., Fadel, H.J., Lee, K., et al. (2020). CPSF6-Dependent Targeting of Speckle-Associated Domains Distinguishes Primate from Nonprimate Lentiviral Integration. mBio 11. 10.1128/mBio.02254-20.

