## Supplementary material for "The role of LEDGF in transcription is exploited by HIV-1 to position integration": Suppl. Figures

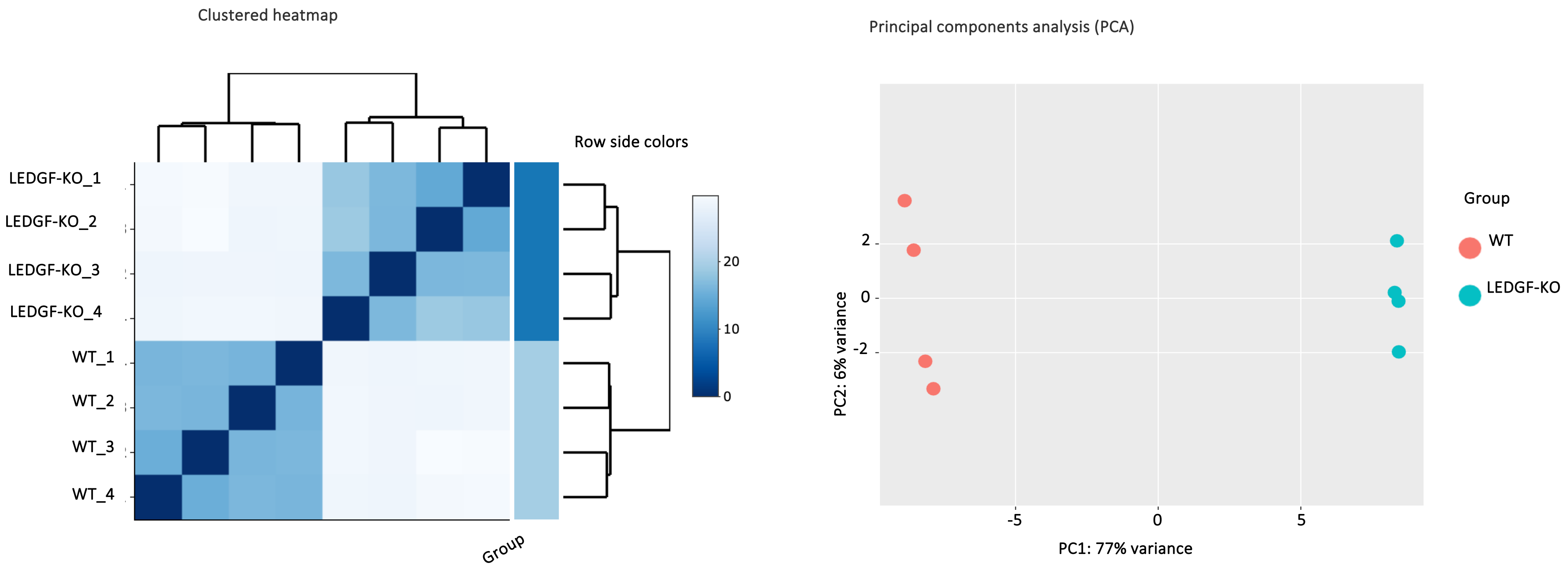


Suppl. Fig. S1. Left panel. The heatmap shows a hierarchical clustering of pairwise distances between samples. Darker blue means less distant (i.e. more similar). In general we expect to see replicates clustering together and separation of treatments. Right panel. Principal components analysis (PCA). The x- and y-axes do not have units, rather, they represent the dimensions along which the samples vary the most. The amount of variance explained by each principal component is indicated in the axes label.

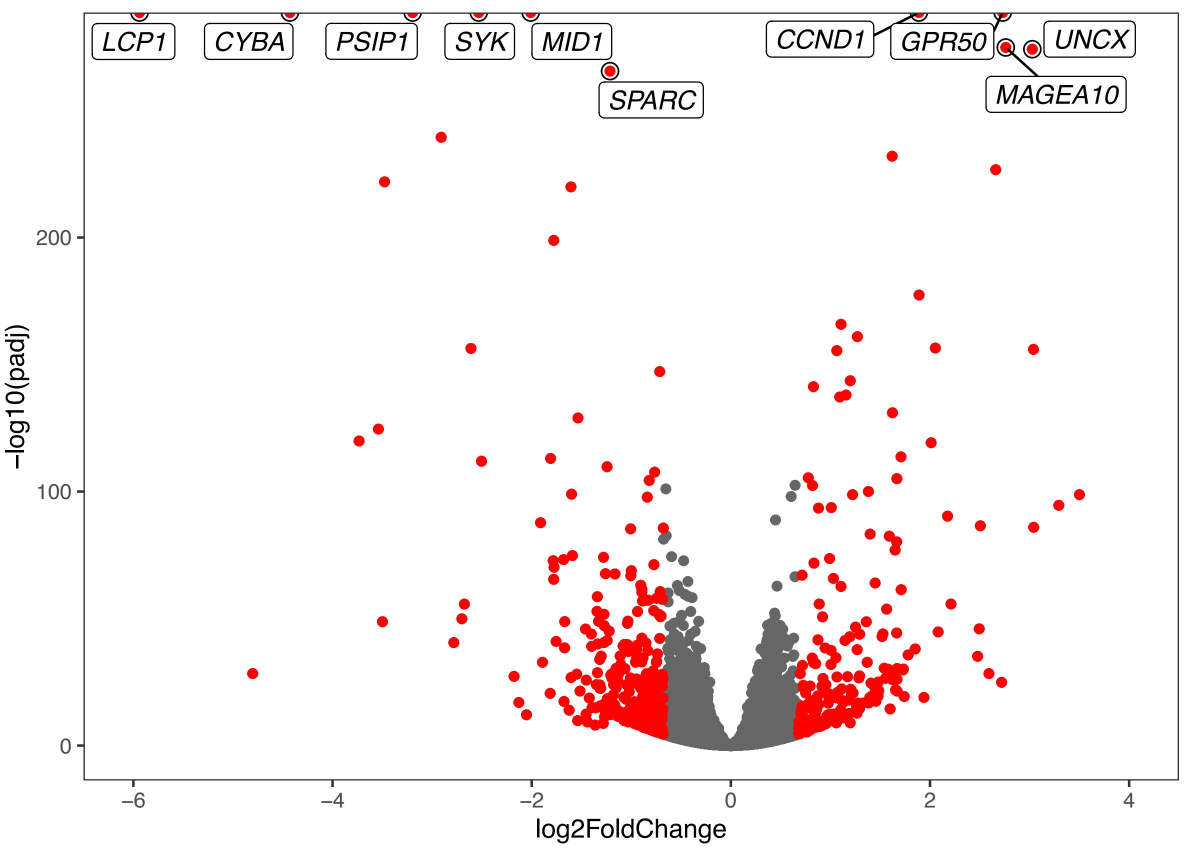


Suppl. Fig. S2. Volcano plot where the differentially expressed genes of HEK293T WT and LEDGF KO (padj < 0.1) with an absolute fold-change > 1.6 (or LFC > log2(1.6)) are color-coded in red.

MID1, Differential TSS isoforms

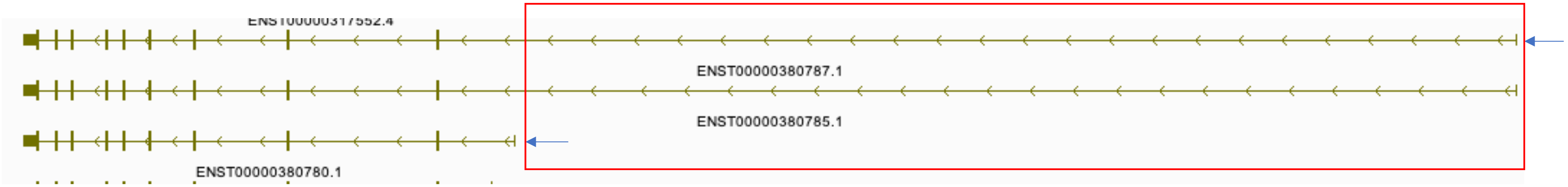


NUCB2, Differential TSS and Splicing isoforms

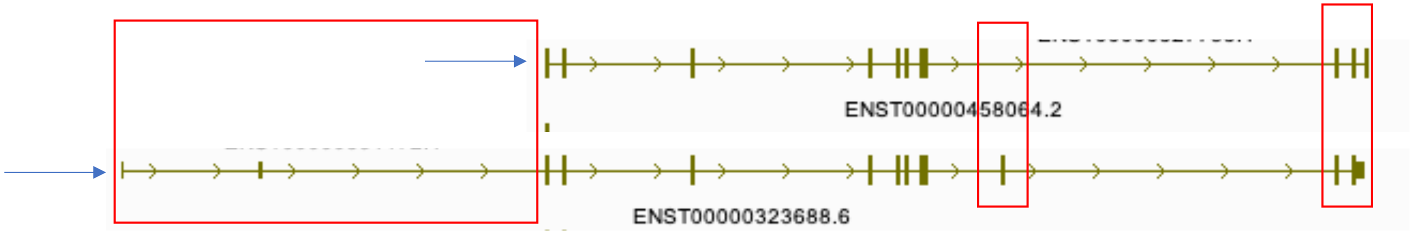


FBXW11, differential splicing isoforms

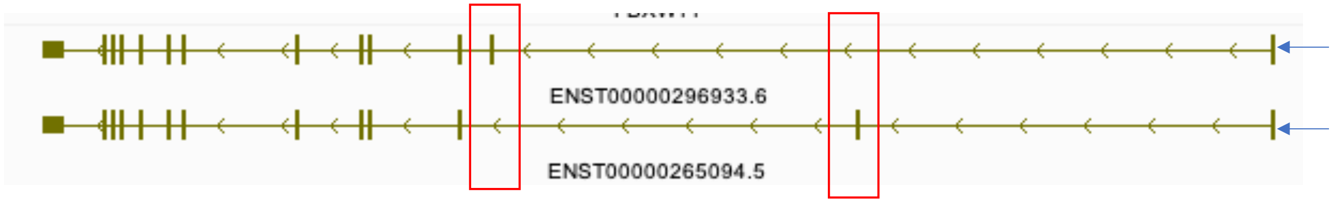


Suppl. Fig. S3. Examples of differential usage transcript isoforms of WT vs LEDGF KO cells exhibiting different TSS and splice choices.

H3K4me3

RNA Pol II

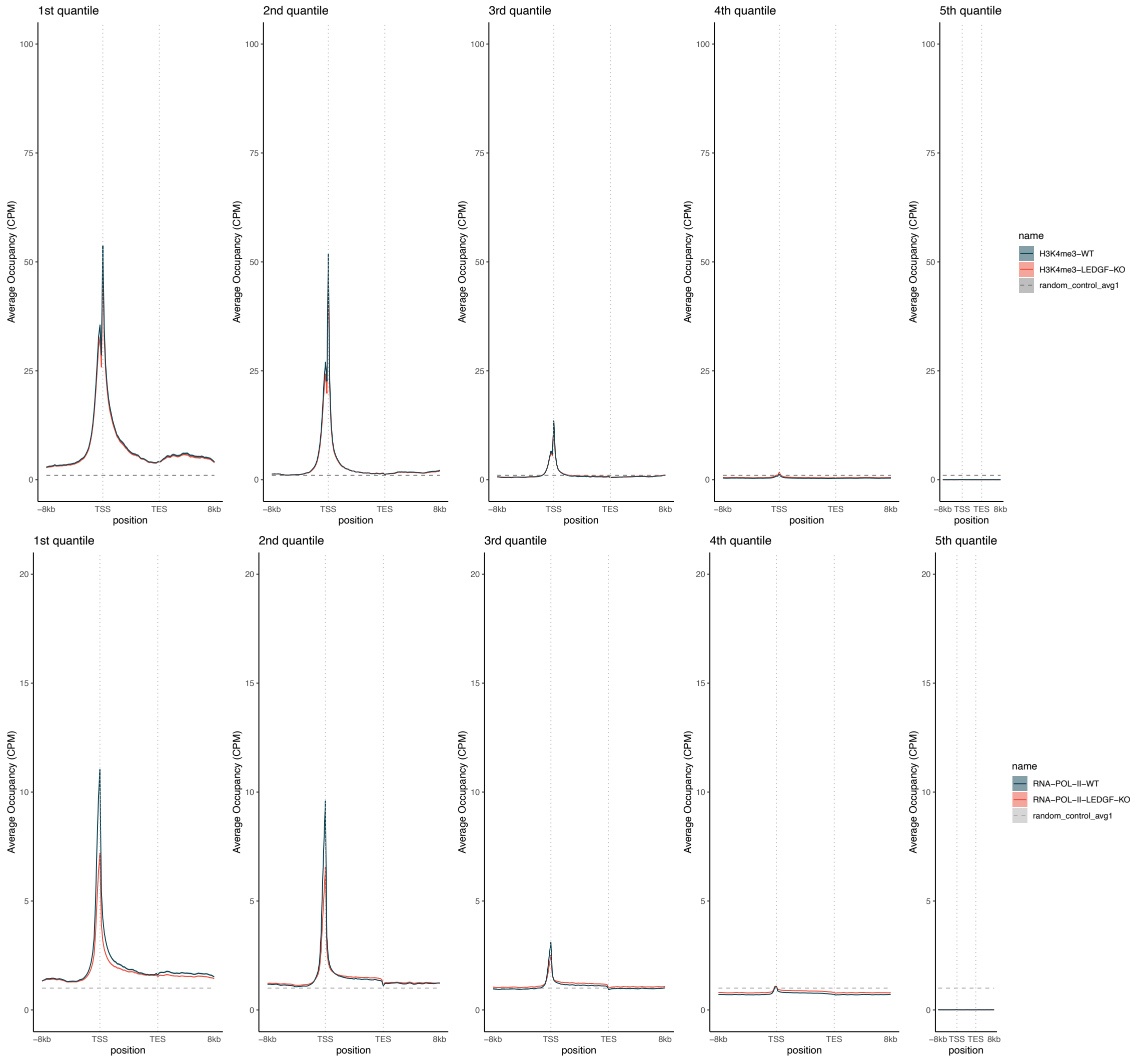


Suppl. Fig. S4. Metagene plots of H3K4me3 and RNA Pol II of genes placed in five quantiles sorted by levels of LEDGF-3XFLAG. Black plots are from WT cells and red plots are from LEDGF KO cells.

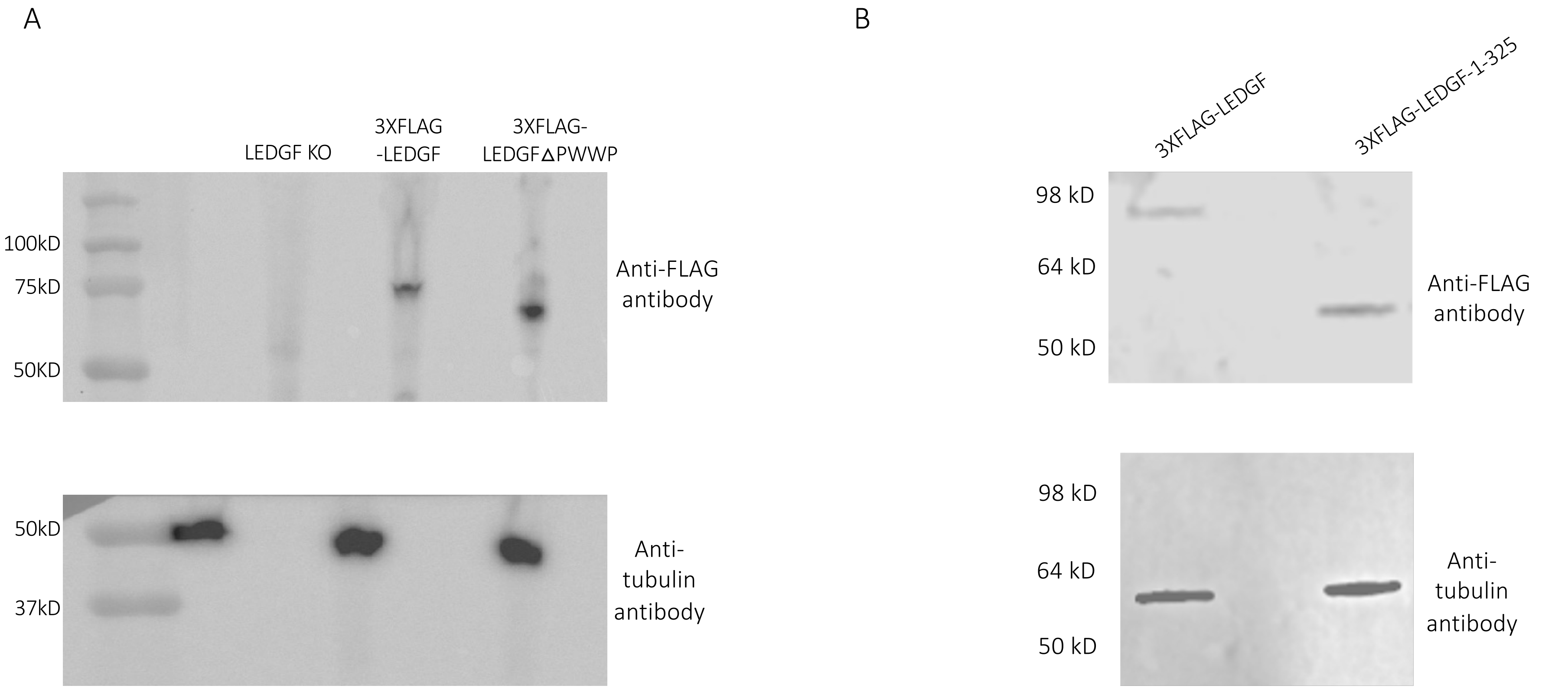


Suppl. Fig. S5. A. Immunoblot of HEK293T LEDGF KO, KO expressing 3XFLAG-LEDGF, or 3XFLAG-LEDGFΔPWWP. B. Immunoblot of HEK293T LEDGF KO expressing 3X-FLAG-LEDGF or 3X-FLAG-1-325.

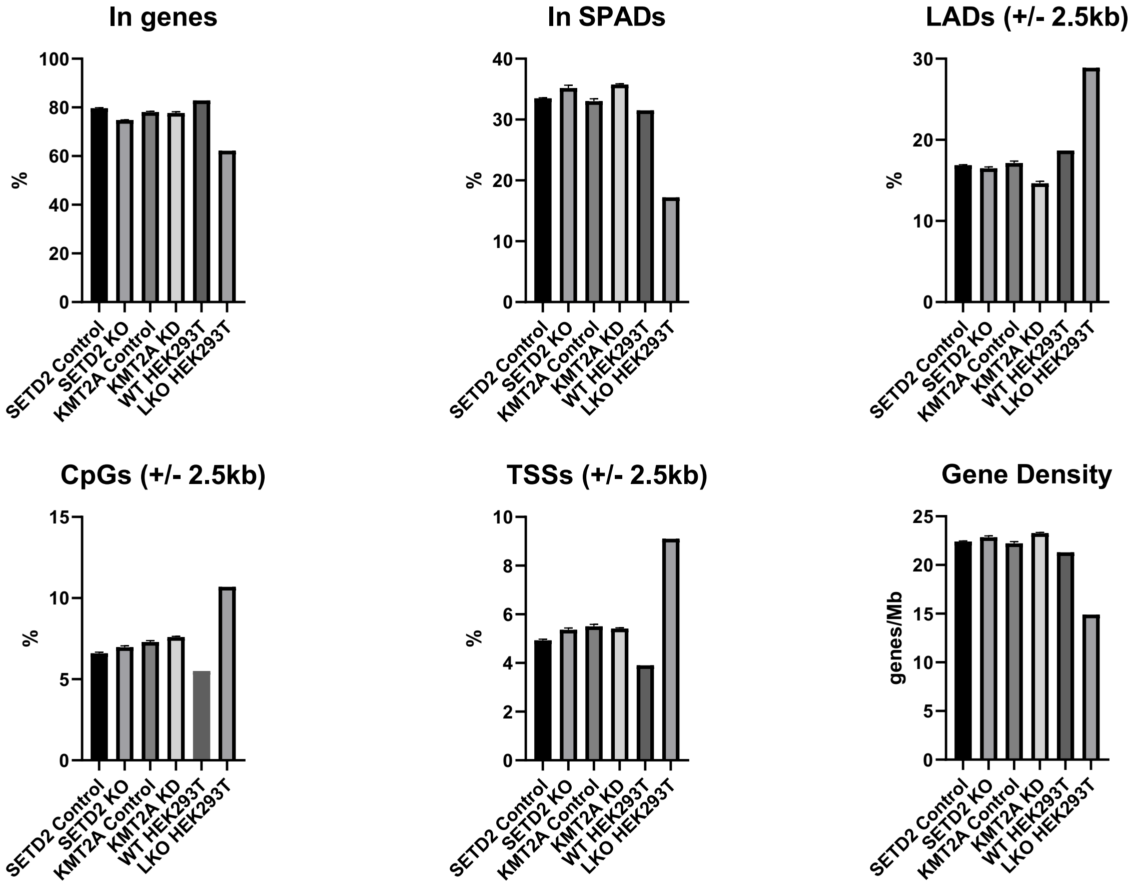


Suppl. Fig. S6. Frequencies of HIV-1 integration within the indicated portions of the human genome for the indicated knockout/knockdown and matched control samples. Historical WT HEK293T and LEDGF KO (LKO) cell data are from Li W et al., 2020 mBio, Vol 11:e02254-20.

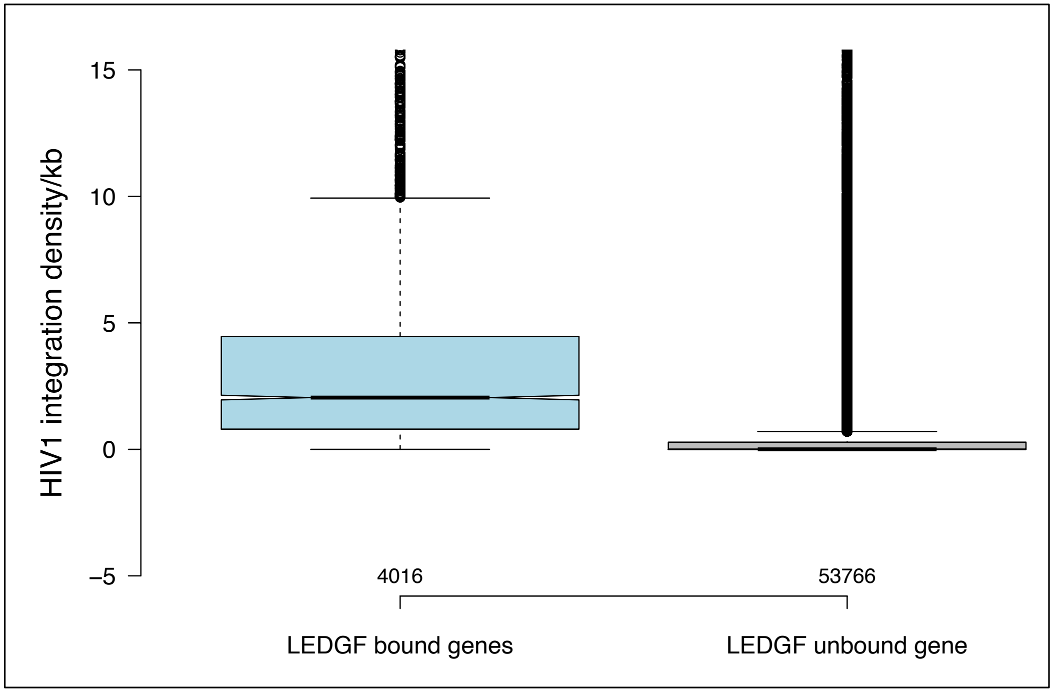


Suppl. Fig. S7. Average integration density in genes with >2.5-fold enrichment of LEDGF-3XFLAG and genes with less enrichment of LEDGF-3XFLAG.

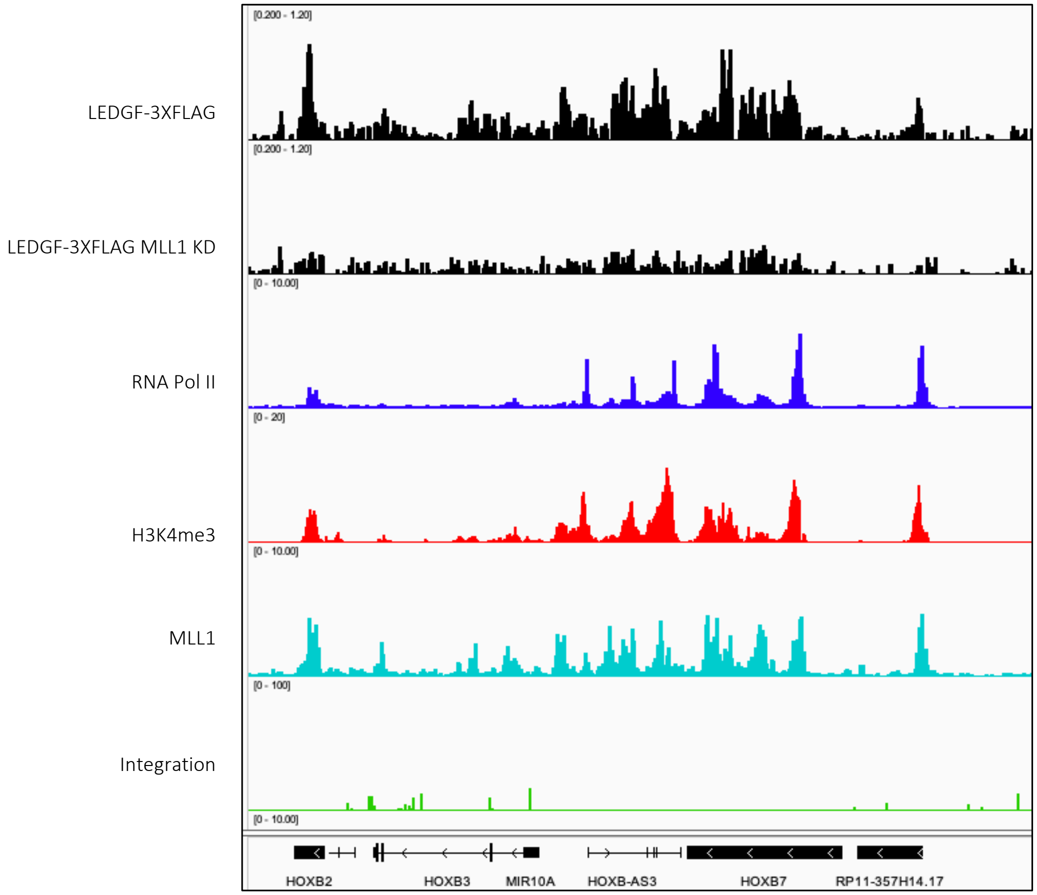


Suppl. Fig. S8. HOX gene cluster with enrichment of LEDGF-3XFLAG, LEDGF-3XFLAG in MLL1 KD, MLL1, RNA Pol II, H3K4me3 and with very little integration. LEDGF-3XFLAG association is dependent on MLL1, suggesting IBD bound by MLL1.

Immunoblotting of h7,h8, h9 and h10 HEK-293 T cell lysates on the day of infection

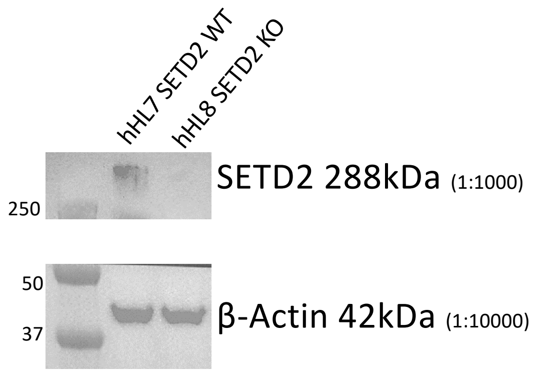


Suppl. Fig. S9. Immunoblot of HEK293T cells with WT SETD2 and SETD2 KO. Antibodies were specific for SETD2 and β-Actin.
